# CA1 pyramidal cell diversity is rooted in the time of neurogenesis

**DOI:** 10.1101/2021.03.07.433031

**Authors:** Davide Cavalieri, Alexandra Angelova, Anas Islah, Catherine Lopez, Marco Bocchio, Agnès Baude, Rosa Cossart

## Abstract

Cellular diversity supports the computational capacity and flexibility of cortical circuits. Accordingly, principal neurons at the CA1 output node of the hippocampus are increasingly recognized as a heterogeneous population. Their genes, molecular content, intrinsic morpho-physiology, connectivity, and function seem to segregate along the main anatomical axes of the hippocampus. Since these axes reflect the temporal order of principal cell neurogenesis, we directly examined the relationship between birthdate and CA1 pyramidal neuron diversity, focusing on the ventral hippocampus. We used a genetic fate-mapping approach that allowed tagging three groups of age-matched principal neurons: pioneer, early- and late-born. Using a combination of neuroanatomy, slice physiology, connectivity tracing and cFos staining, we show that birthdate is a strong predictor of CA1 principal cell diversity. We unravel a subpopulation of pioneer neurons recruited in familiar environments with remarkable positioning, morpho-physiological features, and connectivity. Therefore, despite the expected plasticity of hippocampal circuits, given their role in learning and memory, the diversity of their main components is significantly predetermined at the earliest steps of development.

## Introduction

Hippocampal circuits serve multiple complex cognitive functions including navigation, learning, and episodic memory. For this purpose, they form highly associative networks integrating external inputs conveying multi-sensory, proprioceptive, contextual and emotional information onto internally-generated dynamics. For many years, in contrast to their inhibitory counterparts, principal glutamatergic cells have been treated experimentally and modeled computationally as identical twins. Hence, multiple streams of information serving a variety of functions were considered to be integrated by a uniform group of neurons. This conundrum has been recently clarified by converging results indicating that the hippocampus is in reality comprised of heterogeneous principal cell populations forming at least two distinct, nonuniform parallel circuit modules that are independently controlled and involved in different behaviors. This diversity would contribute to the computational flexibility and capacity of the hippocampal circuit (***Soltesz and Losonczy, 2018; Valero and de la Prida, 2018***).

At population level, it is becoming increasingly evident that most features of neuronal diversity within the CA1 output node of the hippocampus distribute in a way that matches the temporal schedules of development across transverse and radial axes. According to early autoradiographic studies, mouse CA1 principal neurons (CA1 PNs) are born between E12 and birth with a peak at E14 (***Angevine, 1965; Bayer, 1980***). In the radial axis, successive generations of PNs occupying the principal pyramidal layer migrate past the existing earlier born neurons thus creating layers in an “inside-out” fashion. Therefore, superficial neurons (closer to the *stratum radiatum*) are in general born later than deep neurons (closer to the *stratum oriens*). In the transverse axis, distal CA1 neurons (CA1a and b, closer to the subiculum) are born first, followed by proximal CA1 (closer to CA2). In contrast, there is no apparent developmental gradient along the dorsoventral axis in CA1 as compared to CA3 or entorhinal cortex, where ventral neurons are born significantly later that dorsal ones (***Bayer, 1980; Donato et al., 2017***). In sum, in CA1, early born neurons are preferentially found throughout the dorsoventral axis, in CA1a,b (distal) and in deep radial positions, whereas later born ones are located in CA1c and closer to the *stratum radiatum*.

Heterogeneity of the morpho-physiology of CA1 PNs distributes in a similar pattern. Hence, adult CA1 PNs located in regions associated with a presumed earlier birth date (i.e. CA1 deep and/or distal) display a smaller HCN-mediated h-current (Ih) (***Jarsky et al., 2008; Lee et al., 2014; Maroso et al., 2016; Masurkar et al., 2020***), a lower membrane resistance (Rm) (***Graves et al., 2012; Masurkar et al., 2020***) as well as a higher excitability (***Cembrowski et al., 2016; Mizuseki et al., 2011***) and bursting propensity (***Jarsky et al., 2008***). The same applies to the local and long-range connectivity schemes of CA1 PNs. For example, seminal early studies noted how the order of neurogenesis in the entorhinal cortex, proceeding from lateral to medial, also strictly correlated with the order of its termination on CA1 PNs along transverse and radial axes (***Bayer, 1980***); a property recently probed at functional level (***Masurkar et al., 2017***). The temporal order of neurogenesis was directly evidenced at single-cell level to impose the local patterning of glutamatergic connectivity to form isochronic circuits throughout the hippocampal trisynaptic circuit (***Deguchi et al., 2011***). This also applies for the mesoscopic organization of adult GABAergic inhibitory circuits. Indeed, both along the radial and transverse axes of development, late born PNs and subregions are more likely to drive CA1 interneurons, while early born regions and PNs receive stronger inhibitory inputs (***Donato et al., 2017; Lee et al., 2014; Oliva et al., 2016; Valero et al., 2015***).

Finally, functional diversity also appears to correlate with presumed birthdate. When combining the information of many recent reports a picture emerges by which in the dorsal hippocampus putatively early born PNs are comprised of a higher fraction of place-modulated neurons (***Danielson et al., 2016; Mizuseki et al., 2011***) with a poorer spatial coding specificity (***Danielson et al., 2016; Geiller et al., 2017; Hartzell et al., 2013; Henriksen et al., 2010; Oliva et al., 2016***). More particularly, in CA1, presumably older PNs are better tuned to receive external sensory inputs as their firing is more anchored to external landmarks while later born ones, may be more likely to convey an internal “memory stream”, more likely to participate in SWRs and to convey self-referenced information, with slower if any remapping and more stable place maps (***Danielson et al., 2016; Fattahi et al., 2018; Sharif et al., 2020; Mizuseki et al., 2011; Valero et al., 2015***). Similarly, function segregates along the radial axis in the ventral hippocampus where deep CA1PNs specifically projecting to the nucleus accumbens shell (NAcc) or to the Lateral hypothalamus (LHA) were reported to contribute to social and anxiety-related behaviors, respectively (***Jimenez et al., 2018; Okuyama et al., 2016***).

Altogether, mostly based on the tight correlation between their soma position and their morphofunctional attributes, these recent results indirectly support the intriguing possibility that CA1PNs diversity may be partly determined at their time of neurogenesis. Alternatively, diversity may simply reflect final soma position and the influence of local circuits rather than an early predetermination. In order to test directly these two non-mutually exclusive possibilities, we have fate-mapped three groups of age-matched CA1PNs: pioneer (i.e. born around embryonic day 12: E12.5), early (E14.5)- and late (E16.5)-born as described previously (***Marissal et al., 2012; Save et al., 2019***). We analyzed their morphophysiological properties, connectivity, and activation during the free exploration of differently familiar environments. We focused on the ventral hippocampus as this region displays the wider diversity of CA1PNs in terms of projection patterns (***Cembrowski et al., 2016; Jimenez et al., 2018; Jin and Maren, 2015; Kim and Cho, 2017; Lee et al., 2014; Okuyama et al., 2016; Parfitt et al., 2017; Xu et al., 2016***).

Whereas the radial position of CA1PNs correlates with some features of synaptic connectivity like the frequency of excitatory synaptic currents they receive, other morphophysiological properties including apical dendritic length in the *stratum radiatum*, parvalbumin somatic coverage, long-range projection, spiking in response to current injections, sag current or input resistance distinguish between cells with different birthdates. In particular, pioneer E12-CA1PNs stand out as a singular subset of cells regarding many parameters, broadly distributed along the radial axis and preferentially activated when mice explore a familiar environment. Therefore, the present study reveals how the heterogeneity of CA1PN diversity extends beyond the mere subdivision into two sequentially generated sublayers along the radial axis. It likely encompasses many, at least three, intermingled subtypes specified at progenitor stage by their temporal origin.

## Results

### Pioneer CA1 pyramidal cells broadly integrate the pyramidal layer and display remarkable features

CA1 pyramidal neurons (PNs) were fate-mapped using the inducible transgenic driver line Ngn2-Cre^ER^, expressing Cre^ER^ under the control of the Ngn2 promoter. We crossed Ngn2-Cre^ER^ mice with the Ai14 reporter line (***Madisen et al., 2010***), including a Cre-dependent TdTomato allele (See Methods and Materials). Like in our previous studies (***Marissal et al., 2012; Save et al., 2019***), three different groups of pyramidal neurons were labelled by TdTomato expression via tamoxifen administration at separate embryonic time points: embryonic day 12.5 (E12.5), E14.5 and E16.5. We first focused on the main features classically expected to segregate into two CA1 sublayers, the *deep* and *superficial* one.

We first examined the somatic location of CA1PNs in the *stratum pyramidale* according to their fate-mapped date of birth Fig. 1A. The location was calculated as the distance between the center of the soma and a line representing the *stratum radiatum/pyramidale* border. Coherently with an inside first-outside last patterning (***Angevine, 1965; Bayer, 1980***), we found that E16.5 CA1PNs were mostly located in the superficial part of the pyramidal layer, whereas the deep sublayer was enriched in E14.5 CA1PNs. In both dorsal and ventral CA1, E12.5 cells were also positioned in the deeper portion, although significantly shifted towards the *stratum oriens*. Noteworthy, not only did the location in respect to the *radiatum/pyramidale* border change, also the spatial dispersion of cells in the pyramidal layer varied with the birthdate. More specifically, E12.5 were the most spread out (E12.5 interquartile range or IQR_E12.5_ = 63.14 in dorsal and 104.44 in ventral CA1) and later born cells showed drastically reduced values of dispersion (IQR_E14.5_ =22 and 30.71, IQR_E16.5_ = 22 and 12.95 dorsally and ventrally, respectively). The broad distribution of E12.5PNs and its shift towards the *stratum oriens* was particularly obvious in cumulative distribution plots obtained from dorsal and ventral CA1 where a consistent fraction of the labelled E12PNs were located outside from the *stratum pyramidale*, in the *stratum oriens* and, fewer, in the *stratum radiatum* Fig. 1B. These results are globally in line with previous work. However, they indicate how not only the location, but also the dispersion of somata in the pyramidal layer depends on the birth date. In addition, they show how the deep CA1 sublayer is comprised of a mixed population of E12.5 and E14.5 CA1PNs while cells located in the *stratum oriens* are predominantly of E12.5 origin.

**Figure 1.**
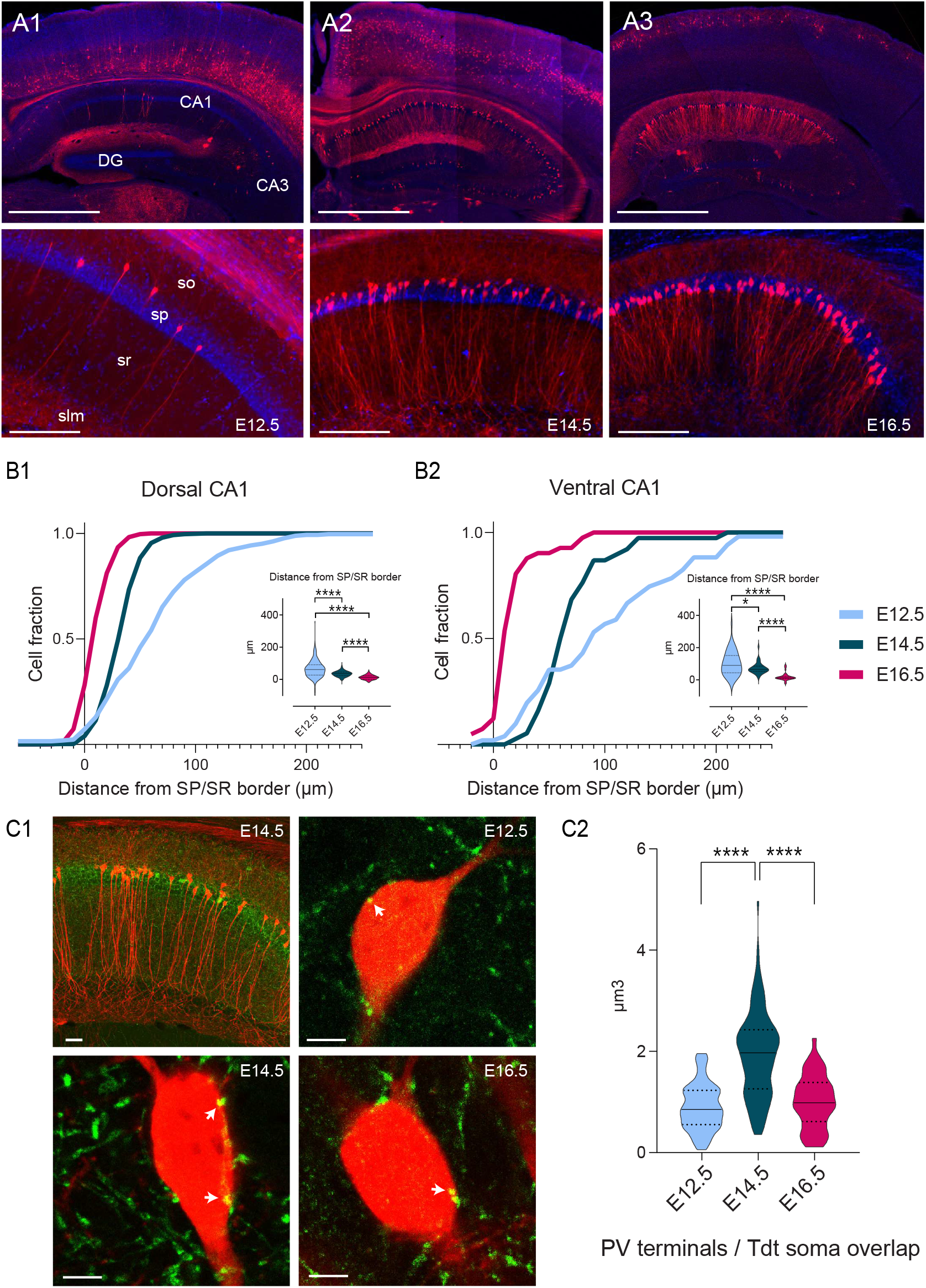
Soma location distribution and PV innervation of CA1PNs changes according to birthdate (A1-3). Top, representative sections of the dorsal hippocampus and cortex, illustrating Tdtomato (Tdt) labelling in CA1 pyramidal neurons (PNs) in Ngn2-Cre^ER^-Ai14 mice after tamoxifen induction at embryonic day 12.5 (E12.5), E14.5 and E16.5, from left to right. Bottom, higher magnification on CA1. E12.5 PNs are rare and dispersed, while E14.5 and E16.5 are predominantly found in the deep (upper) and superficial (lower) portion of the pyramidal layer, respectively. Note that in the above cortical areas, E12.5 PNs are restricted to the deepest layers, E14.5 PNs the middle ones (IV-III) and E16.5 PNs are most superficial (layers I-II). Scalebars: top, 1000 µm; bottom, 200 µm. Cx: Cortex; DG: dentate gyrus; so: stratum oriens; sp: stratum pyramidale; sr: stratum radiatum; slm: stratum lacunosum-moleculare. **B1-2** Quantification of the soma location distribution of birth-dated CA1 PNs. **B1** Cumulative fraction of E12.5, E14.5 and E16.5 PNs in dorsal CA1, calculated as distance in µm from the superficial (lower) border of the stratum pyramidale. Insert, E12.5PNs are located further away from the border than E14.5PNs (P<0.0001, CI_95%_ [18.18; 33]) and E16.5 PNs (P<0.0001, CI_95%_ [41; 55.73]). In turn, E14.5PNs also occupy deeper positions than E16.5 PNs (P<0.0001, CI_95%_ [21; 25]). **B1** Cumulative fraction of E12.5, E14.5 and E16.5 PNs in ventral CA1. Insert, similarly to dorsal CA1, E12.5PNs are located deeper than E14.5 PNs (P: 0.0175, CI_95%_ [3.13; 57.72]) and E16.5 PNs (P<0.0001, CI_95%_ [53.93; 109.57]), and E14.5 PNs than E16.5PNs (P<0.0001, CI_95%_ [44.47; 59.32]). **C1-2** Parvalbumin (PV) innervation onto birth-dated CA1 PNs. **C1** Top left, representative view of the CA1 area in a Ngn2-CreER-Ai14 mouse tamoxifen-induced at E14.5, with Tdt^+^ cells (red) and PV cells (green). Top right, close-up on the soma of E12.5 PNs (red) displaying putative somatic PV boutons (indicated by a white arrow). Bottom left, close-up on E14.5 PNs. Bottom right, close-up on E16.5 PNs. **C2** Quantification of the volume overlap (µm) of putative PV terminals with Tdt^+^ soma depending on the birthdate. E14.5 PNs present more putative boutons than E12.4 (P<0.0001, CI_95%_ [-1.264; -0.81]) and E16.5 (P<0.0001, CI_95%_ [0.66; 1.22]), while the latter two are not significantly different (P: 0.7038, CI_95%_ [-0.389; 0.19]). Scalebars: top left: 50 µm; top right, bottom left and bottom right: 5 µm.

We next investigated the diversity among CA1PNs with parvalbumin, a classical immunohistochemical marker used in several previous studies to distinguish the deep and superficial sublayers. Indeed, parvalbumin was demonstrated to preferentially decorate the somata of deep CA1PNs, indicating a stronger innervation by putative PV-basket cells (***Lee et al., 2014; Soltesz and Losonczy, 2018; Valero et al., 2015***). For this reason, we expected to find a preferential PV innervation of E12.5 and E14.5 PNs, both located in the deep sublayer Fig. 1C. Surprisingly, this labelling was better segregating according to birthdate than position. Indeed, E12.5CA1PNs displayed significantly less putative PV contacts than E14.5CA1PNs, even though their somata were mainly located in the deep CA1 (see Fig. 1B). These first observations indicate that, at single-cell level, birthdate may better correlate with the properties of CA1PNs than position along the radial axis. They also point-out at the pioneer subset of CA1PNs as potential outliers in the relationship between cell diversity and birth order.

### Synaptic input drive onto CA1PNs reflect both radial positioning and birthdate

To further analyze the relationship between birthdate, cell position and single-cell morpho-physiological features, we next performed a series of ex vivo whole cell patch clamp recordings in acute brain slices from adult Ngn2-Cre^ER^-Tdt mice (n = 72 cells from 34 mice; 22 E12.5 CA1PNs, 27 E14.5 CA1PNs, 23 E16.5 CA1PNs), sampling from intermediate to ventral CA1.

Previous studies suggest that cells located in the deep or superficial portion of the CA1 pyramidal layer, differ in their synaptic inputs (***Kohara et al., 2014; Lee et al., 2014; Masurkar et al., 2017; Valero et al., 2015***). In order to investigate whether this pattern would also depend on birthdate we have voltage-clamped the fate-mapped CA1PNs at the reversal potential for GABAergic and glutamatergic currents to record spontaneous excitatory and inhibitory postsynaptic currents (s-EPSCs and s-IPSCs, Fig. 2). The location of the neurobiotin-filled somata of the recorded cells was measured in respect to the depth of the pyramidal layer and expressed as a ratio between 0 and 1, representing the border with the *stratum radiatum* and the *stratum oriens* respectively. Although the amplitude and frequency of neither s-EPSCs (Fig. 2C) nor s-IPSCs (Fig. 2D) differed, the excitatory/inhibitory balance (computed as the E/I amplitude ratio, Fig. 2E) was lower for E14.5 than both E12.5 (P: 0.020) and E16.5 CA1PNs (P: 0.018). This indicates that neurons born at E14.5 are subject to a synaptic drive leaning more towards inhibition that the other two groups, in agreement with their receiving more putative perisomatic PV contacts (see above). In contrast, the frequency of s-EPSCs (Fig. 2C) showed a linear correlation with the cell body location (r= 0.49, p: 0.002). This was also reflected in the E/I frequency ratio (r=0.43, P: 0.0047), indicating a higher excitatory synaptic drive in deep than in superficial neurons, regardless of their birthdate (Fig. 2E).

**Figure 2.**
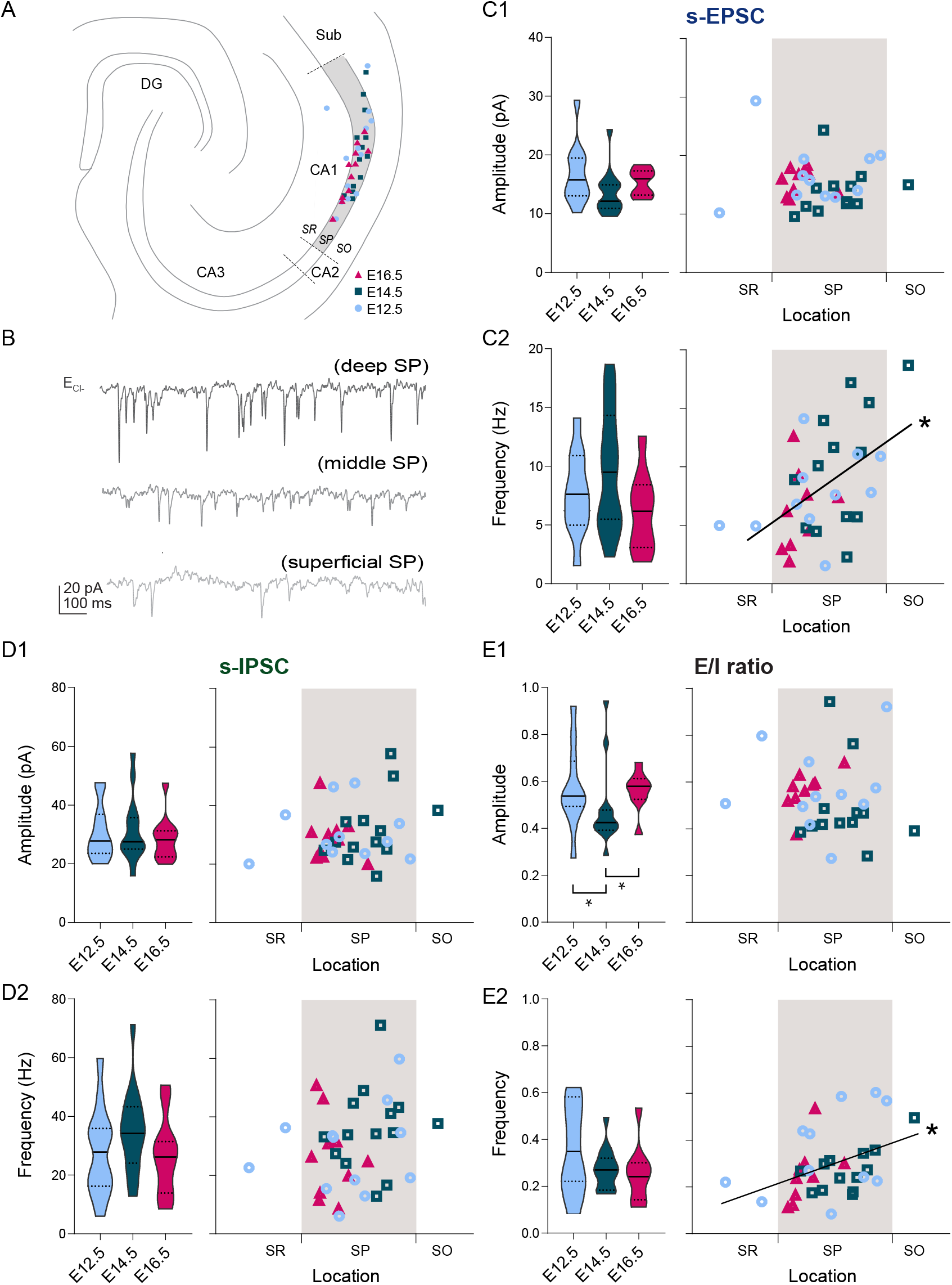
Overall synaptic drive is defined by the time of neurogenesis. **(A)** Anatomical location of neurobiotin-filled CA1PNs recorded from acute horizontal slices in voltage-clamp experiments. DG: dentate gyrus, Sub: subiculum, SR: stratum radiatum, SP: stratum pyramidale, SO: stratum oriens. **(B)** Representative traces of spontaneous excitatory synaptic currents (s-EPSCs) recorded cells at chloride ion reversal potential (E_Cl_-) from three CA1PNs, located respectively in the deep, middle, and superficial portion of the pyramidal layer (SP). Note that the occurrence of synaptic events decreases from top (deep) to bottom (superficial). **(C1)** s-EPSC amplitude recorded in E12.5, E14.5 and E16.5 CA1PNs. On the left panel, violin plot of E12.5, E14.5 and E16.5 PNs s-EPSC amplitude. On the right panel, color-coded scatterplot of the same data plotted against the radial position of each cell. No linear correlation between s-EPSC amplitude and location was found. **(C2)** s-EPSC frequency recorded in E12.5, E14.5 and E16.5 CA1PNs. On the left panel, violin plot of the three birthdate groups. On the right panel, scatterplot of the same data against the radial position. S-EPSC frequency increases linearly with the cell location, the two being positively correlated (P: 0.017; CI_95%_ [0.186; 0.710]). The deeper, the more frequent spontaneous excitatory events a cell receives. **(D1)** s-IPSC amplitude recorded in E12.5, E14.5 and E16.5 CA1PNs. On the left panel, violin plot of the three birthdate groups. On the right panel, scatterplot of the same data against the radial position. No linear correlation between s-IPSC amplitude and location was found. **(D2)** s-IPSC frequency recorded in E12.5, E14.5 and E16.5 CA1PNs. On the left panel, violin plot of the three birthdate groups. On the right panel, scatterplot of the same data against the radial position. No linear correlation between s-IPSC frequency and location was found. **(E1)** Ratio between EPSC and IPSC (E/I) amplitude recorded in E12.5, E14.5 and E16.5 CA1PNs. On the left panel, violin plot of the three birthdate groups. E14.5 PNs display a lower ratio than E12.5 PNs (P_adj_: 0.010; CI_95%_ [0.031; 0.210]) and E16.5 PNs (P_adj_: 0.006; CI_95%_ [-0.193; -0.057]), suggesting that their overall synaptic drive leans more towards inhibition than the other two groups. On the right panel, scatterplot of the same data against the radial position. No linear correlation between E/I amplitude ratio and location was found. **(E2)** E/I frequency ratio recorded in E12.5, E14.5 and E16.5 CA1PNs. On the left panel, violin plot of the three birthdate groups. On the right panel, scatterplot of the same data against the radial position. E/I ratio linearly correlates with the cell location, similarly to s-EPSC frequency (P: 0.0043; CI_95%_ [0.126; 0.654]). Violin present medians (center), interquartile ranges (bounds), minima and maxima. Color-code: E12.5: light blue, E14.5: dark blue, E16.5: magenta. The gray shaded area in scatterplots represents the thickness of the stratum pyramidale. *P < 0.05.

To acquire further understanding of the different connectivity profiles, we applied brief electric stimulations in the stratum radiatum, which contains CA3 Schaffer collaterals innervating CA1, and recorded evoked PSCs in voltage-clamp mode (Fig. 3A). In contrast to (***Valero et al., 2015***), we could not observe any correlation between the properties of evoked synaptic excitation and the soma location or the birth date (Fig. 3B & Fig. 3C). This contrasted with evoked inhibitory responses (Fig. 3B & Fig. 3D). Indeed, evoked IPSC amplitude increased linearly towards the *stratum oriens* (r=0.34, P: 0.0128), while IPSC kinetics (half-width and time constant) inversely correlated with the location (respectively, r=-0.36, P: 0.008 and r=-0.36, P: 0.0051, Fig. 3D). This suggests that cells in the deep sublayer receive stronger and faster presumably CA2/CA3-driven GABAergic input (***Lee et al., 2014***). Finally, we tested paired-pulse (PP) responses as a proxy of the input release probability and found that inhibitory PP ratio was consistently higher in E12.5 CA1PNs as compared to E14.5 (P: 0.0326) and E16.5 (P: 0.0177) CA1PNs (Fig. 3D).

**Figure 3.**
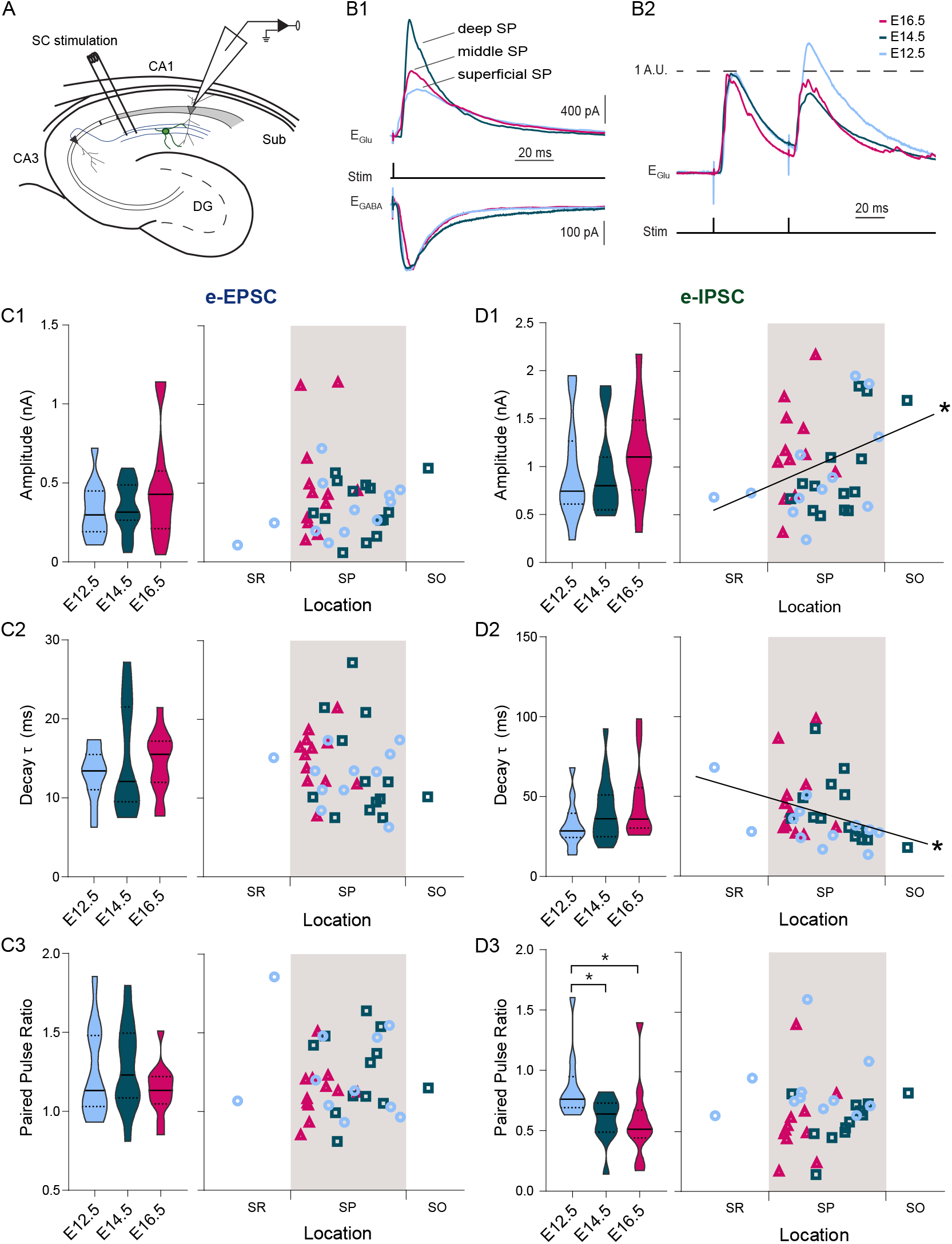
Both developmental and positional factors determine Schaffer collateral-evoked inhibition. **(A)** Experimental paradigm in which synaptic responses to Schaffer collaterals (SC) stimulation are recorded from CA1PNs. DG: dentate gyrus, Sub: subiculum. **(B1)** Representative mean traces of evoked inhibitory (top) and excitatory (bottom) synaptic currents (e-IPSCs and e-EPSCs) recorded upon electric stimulation in the stratum radiatum from CA1PNs, located respectively in the deep, middle, and superficial portion of the pyramidal layer (SP). Note that the amplitude and kinetics of e-IPSCs vary from deep to superficial. **(B2)** Representative mean traces of paired pulse response recorded at glutamatergic receptor reversal potential (E_Glu_) and normalized on the first pulse amplitude. The current response to the second pulse in the E12.5PNS shown is proportionally larger than in E14.5 and E16.5 CA1PNs. **(C1-3)** e-EPSC amplitude (C1), decay (C2) and paired pulse ratio (C3, PPR) recorded in E12.5, E14.5 and E16.5 CA1PNs. Left, violin plot of the three birthdate groups. Right, scatterplot of the same data against the radial position. No linear correlation between s-IPSC amplitude and location was found in any of the three measures. **(D1-D3)** e-IPSC amplitude (D1), decay time constant τ (D2) and PPR (D3) recorded in E12.5, E14.5 and E16.5 CA1PNs. Left, violin plot of the three birthdate groups. Right, scatterplot of the same data against the radial position. e-IPSC amplitude (P: 0.013; CI_95%_ [0.048; 0.576]) and decay τ (P: 0.0051; CI_95%_ [-0.599; -0.097]) display a positive and negative correlation with cell location, respectively. Overall, deeper cells are subject to larger and faster SC-associated inhibitory currents than superficial ones. PPR is higher in E12.5 CA1PNs than in E14.5 (P_adj_: 0.0326; CI_95%_ [0.015.; 0.342]) and E16.5 (P_adj_: 0.0177; CI_95%_ [0.083; 0.452]). Violin present medians (center), interquartile ranges (bounds), minima and maxima. Color-code: E12.5PNs: light blue, E14.5: dark blue, E16.5: magenta. The gray shaded area in scatterplots represents the thickness of the stratum pyramidale. *P < 0.05

Altogether these results indicate that input connectivity motifs might be expressed as a gradient through the depth of the pyramidal cell layer. However, they also show that some functional features, such as the E/I balance and the SC-associated GABAergic release probability, may rather be established through developmental programs that are revealed uniquely by the temporal origin.

### Relationship between birthdate, soma position and synaptic output

We have seen above that the radial position of CA1PNs is partly reflected in their synaptic input drive. Given that these cells were shown to be diverse regarding their projection area, we decided to test whether birthdate could segregate CA1PNs with different projection patterns. The ventral CA1 is known to display multiple projections targeting the EC, the amygdala (Amy), nucleus accumbens (NAcc), the medial prefrontal cortex (mPFC), the lateral septum (LS) and lateral hypothalamus (LHA) (***Arszovszki et al., 2014; Cenquizca and Swanson, 2007; Ciocchi et al., 2015; Kim and Cho, 2017; van Groen and Wyss, 1990***). Interestingly, an anatomical segregation in a laminar fashion was often reported when comparing cells with different projections.

To this aim, we performed injections of the retrograde tracer cholera toxin subunit B (CTB) CTB-647 into the Amy, NAcc, mPFC, LS and LHA and counted CTB^+^-Tdt^+^ co-labelled cells in the ventral CA1 (Fig. 4, and Fig. 4-Supplement1). Co-labelling data from a given animal was translated into a binary vector of length N equal to the total amount of Tdt^+^-birthdated cells, with “1” entries reporting the number of those that were also positive for CTB. Vectors representing animals for which the same birth date was labelled were added. Birth dated groups were next compared in a pairwise-fashion using a resampling approach (See Methods and Materials).

**Figure 4.**
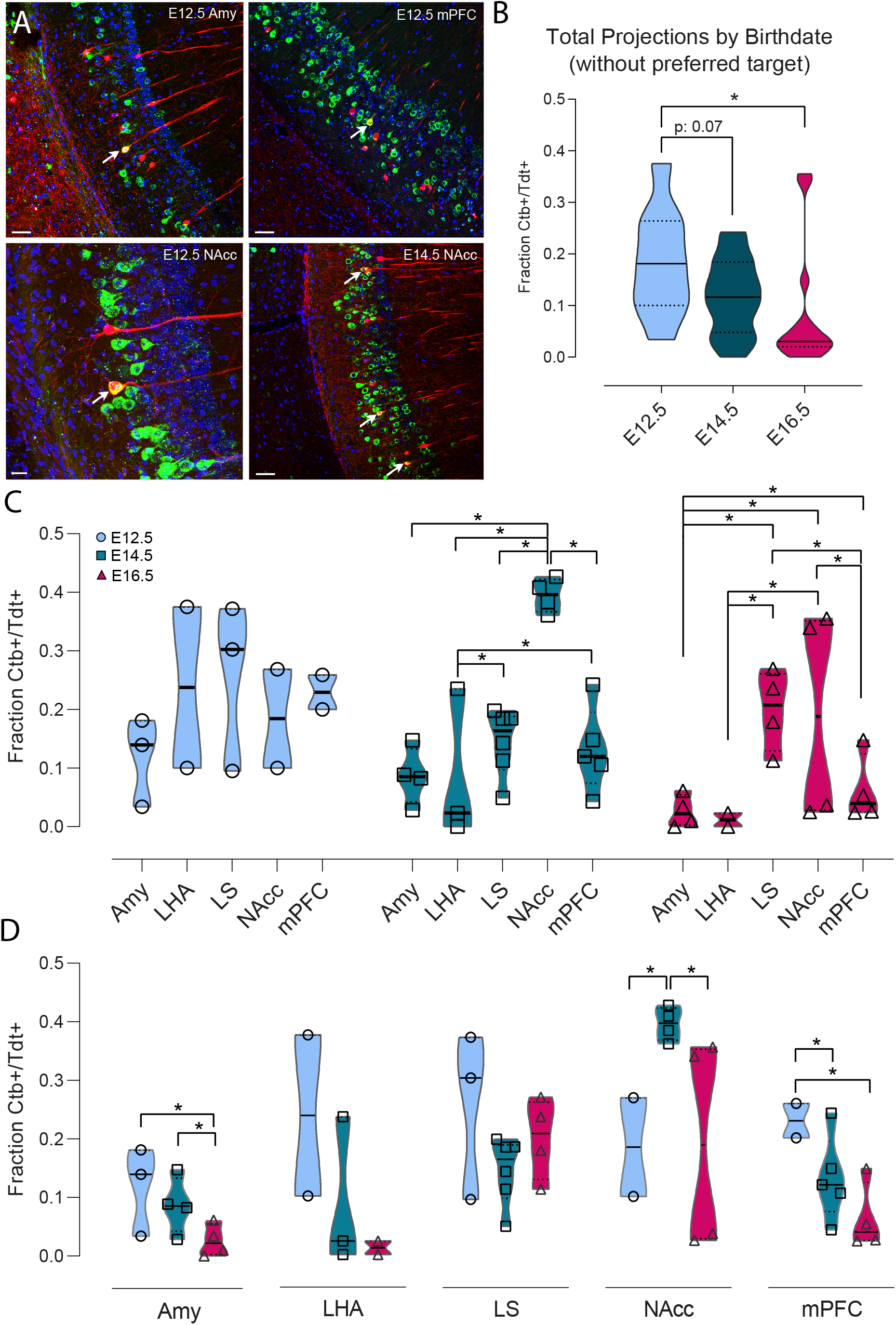
Ventral CA1 PNs with different birthdates exhibit distinct output connectivity. **(A)** Examples of cholera toxin subunit b (Ctb) retrograde labelling (green) in ventral CA1 after injection in amygdala in Ngn2-Cre^ER^-Ai14 mouse induced at E12.5 (top left), medial prefrontal cortex in E12.5 mouse (top right), Nucleus Accumbens (NAcc) in E12.5 mouse (bottom left) and NAcc in E14.5 mouse (bottom right). Colabelled Ctb^+^/Tdt^+^ cells are indicated by a white arrow. Scalebars: top left, top right and bottom right, 50 µm; bottom left, 20 µm. **(B)** Quantification of the total Ctb^+^/Tdt^+^ cell fraction for the three birthdates, excluding the preferred region for each (E12.5 = lateral septum (LS), E14.5 = NAcc, E16.5 = LS). Overall, significantly more Ctb^+^/Tdt^+^ were found in E12.5 PNs than E16.5 (P_adj_: 0.0129, CI_95%_ [0.013; 0.19]) and there is also a trend for E12.5 PNs projecting more densely than E14.5 PNs (Padj: 0.07, CI_95%_ [-0.0050; 0.139]). **(C)** Fraction of Ctb^+^/Tdt^+^ cells in E12.5, E14.5 and E16.5 PNs by target region. Note how E12.5 projects more homogeneously to all structures probed, while marked inter-regional differences appear among E14.5 and E16.5 neurons. **(D)** Fraction of Ctb^+^/Tdt^+^ cells in amygdala (Amy), lateral hypothalamic area (LHA), lateral septum (LS), nucleus accumbens (NAcc) and medial prefrontal cortex (mPFC) by time of neurogenesis. E12.5 PNs project more prominently than other birthdate groups to Amy and mPFC, while NAcc is preferentially targeted by E14.5. **Figure 4–Figure supplement 1**. Representative sections

When comparing the three birthdate groups, excluding the regions where each projected the most densely (LS for E12.5 and E16.5 CA1PNs, NAcc for E14.5 CA1PNs), the fraction of CTB^+^-Tdt^+^ was significantly higher in E12.5 than E16.5 (P<0.05) and presented a trend between E12.5 and E14.5 (P=0.070, Fig. 4B). In addition, E12.5 neurons projected homogeneously to all structures analyzed, while marked differences in output innervation were found within E14.5 and E16.5 neurons (Fig. 4C). E14.5 CA1PNs showed a clear bias towards the NAcc, followed by LS and mPFC. Similarly, and despite some variability, LS and NAcc were also preferred outputs of E16.5 CA1PNs, while the remaining three structures (Amy, mPFC, LHA) presented little to no co-labelling.

The overall higher CTB^+^-Tdt^+^ fraction in E12.5 PNs (Fig. 4B) was found as well when looking within given target structures (Fig. 4D). Indeed, E12.5CA1PNs projected proportionally more than E16.5CA1PNs to Amy, and more than both E14.5 and E16.5CA1PNs to mPFC. In contrast, within NAcc-projecting neurons, CTB^+^-Tdt^+^ proportion was the greatest among E14.5 neurons, as compared to E12.5 and E16.5. In addition, E14.5 cells projected more to Amy than their later-generated counterparts (Fig. 4D). These results are in line with previous findings that NAcc, Amy and mPFC are preferentially targeted by earlier-generated neurons located in the deep sublayer (***Jimenez et al., 2018; Lee et al., 2014; Okuyama et al., 2016***). Furthermore, they point at a specific contribution of the embryonic origin in defining output connectivity motifs (e.g. NAcc-projecting neurons mainly identified among E14.5 CA1PNs). In addition, the tendency of E12.5CA1PNs to contact at consistent rates all retro-traced regions is reminiscent of multiple projecting CA1 cells (***Ciocchi et al., 2015; Gergues et al., 2020***).

### Intrinsic electrophysiological properties of CA1PNs reflect birthdate

Together with synaptic input connectivity, intrinsic electrophysiological properties are known to contribute to the specific activation of CA1PNs, for example in the formation of place fields (***Bittner et al., 2017; Epsztein et al., 2011***). They have been shown to vary across cells (***Dougherty et al., 2012; Graves et al., 2012; Maroso et al., 2016; Masurkar et al., 2020; Mizuseki et al., 2011***). We have therefore examined their relationship with birthdate by performing current-clamp experiments in adult slices where CA1PNs from the three age groups could be identified (Fig. 5).

**Figure 5.**
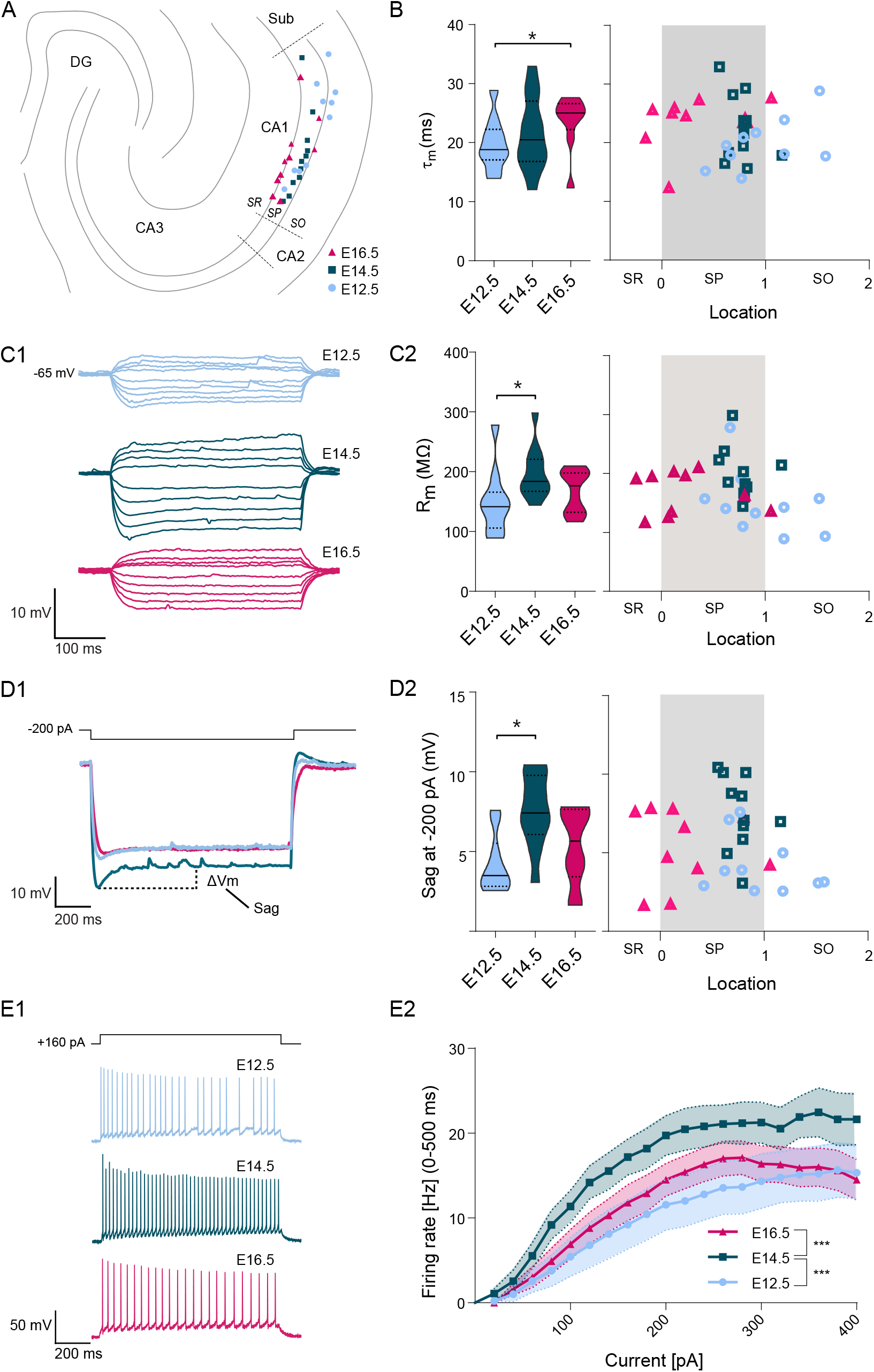
Intrinsic excitability varies according to embryonic birthdate. **(A)** Anatomical location of neurobiotin-filled CA1PNs recorded from acute horizontal slices in current-clamp experiments. DG: dentate gyrus, Sub: subiculum, SR: stratum radiatum, SP: stratum pyramidale, SO: stratum oriens. **(B)** Membrane time constant (τ_m_) of fate mapped CA1PNs. Left, violin plot of the three birthdate groups. Right, scatterplot of the same data against the radial position. E16.5 PNs exhibit a higher τ_m_ than E12.5 cells (P_adj_: 0.0264; CI_95%_ [-8.82; -1.69]). No linear correlation was found with the cell location. **(C1)** Representative membrane potential responses to a series of hyper- and depolarizing current steps recorded from fate mapped CA1PNs. Note the larger deflection of membrane potential in the E14.5 neuron. **(C2)** Membrane input resistance (R_m_) of fate mapped CA1PNs. Left, violin plot of the three birthdate groups. Right, scatterplot of the same data against the radial position. R_m_ is higher in E14.5 than E12.5 cells (P_adj_: 0.0246; CI_95%_ [-89.71; -11.15]). No linear correlation was found with the cell location. **(D1)** Representative depolarizing sag potentials recorded from fate mapped CA1PNs. In this example, E14.5CA1PNs display a greater sag response than E12.5 and E16.5 PNs. **(D2)** Sag potential response of fate mapped CA1PNs, following -200 pA current injections. Left, violin plot of the three birthdate groups. Right, scatterplot of the same data against the radial position. Sag response is significantly higher in E14.5 than E12.5 cells (P_adj_: 0.0026; CI_95%_ [1.74; 6.05]). No linear correlation was found with the cell location. **(E1)** Examples of firing responses to a depolarizing current step recorded from fate mapped CA1PNs. Note that the number of action potential fired by E16.5 and E12.5PNs is lower than E14.5PNs. **(E2)** Input-output curves of fate mapped CA1PNs. E14.5PNs have a higher firing rate than E12.5PNs (P_adj_<0.0001; CI_95%_ [5.21; 11.14]) and E16 (P_adj_<0.0001; CI_95%_ [-9.91; -4.13]), suggesting that they are more excitable. Effect of current injection F (2, 540) = 25.65, P<0.0001; effect of birthdate F (19, 540) = 10.71, P<0.0001; interaction F (38, 540) = 0.2355, P > 0.9999, ordinary two-way ANOVA with Tukey’s post hoc test. Data are represented as means ± standard errors of the means. Violin present medians (center), interquartile ranges (bounds), minima and maxima. Color-code: E12.5: light blue, E14.5: dark blue, E16.5: magenta. The gray shaded area in scatterplots represents the thickness of the stratum pyramidale. *P < 0.05. ***P < 0.001.

We first probed the intrinsic membrane excitability via a series of hyperpolarizing current steps (Fig. 5D) and found that pioneer E12.5CA1PNs exhibited a reduced repolarizing sag response (E12.5 vs E14.5, P: 0.0078), and rebound potential (not shown) in respect to later generated neurons, while their membrane time constant was faster than E16.5 neurons (P: 0.0264, Fig. 5B). On the contrary, pyramidal neurons born at E14.5 had a higher input resistance (Fig. 5C) and, upon depolarizing current injections of increasing amplitude, were globally able to trigger more action potentials than the earlier- and later-born counterparts (Fig. 5E, main effect of birth date P < 0.0001, E12.5 vs E14.5: P < 0.0001, E14.5 vs E16.5: P < 0.0001, E14.5 vs E16.5: P = 0.645, ordinary two-way ANOVA), despite no differences in action potential threshold, half-width or rheobase (see Recapitulative Table).

Interestingly, none of the intrinsic electrophysiological properties measured here correlated with the location in the *stratum pyramidale*, suggesting that the embryonic birth date is a major determinant of the observed cell heterogeneity, which cannot solely be explained by the radial gradient.

### Dendritic morphology of CA1PNs correlates with birthdate

We next asked whether the dendritic morphology of CA1PNs could reflect their birthdate, since pyramidal neurons with different dendritic arborizations were recently shown to distribute radially in the distal part of CA1 (***Li et al., 2017***). We used a set of neurobiotin-filled cells (n_E12.5_ = 30, n_E14.5_ = 14, n_E16.5_ = 19) and considered the overall dendritic arborization (Fig. 6). We measured the distance between the cell soma and the first bifurcation of the primary apical dendrite (Fig. 6B). Earlier-generated neurons (both E12.5- and E14.5-CA1PNs) had a significantly longer shaft than later generated neurons (Fig. 6C, E12.5 vs E16.5, P: 0.0001; E14.5 vs E16.5, P < 0.0001). In addition, the shaft length increased linearly with the distance of the cell body from the *radiatum/pyramidale* border (r=0.51, CI95 [0.32; 0.68], P< 0.0001, Fig. 6C). However, we next reasoned that this correlation could be redundant, owing that a portion of the dendritic shaft includes the distance of the soma from the *radiatum/pyramidale* border, hence its location (Fig. 6B). To avoid this confounding factor, we computed as a substitute the distance between the primary apical dendrite birfucation to the *radiatum/pyramidale* border (Fig. 6C) and observed that indeed the correlation with the soma location could no longer be found (r=0.16, CI95 [-0.07; 0.38], P: 0.0865). However, differences related to the birth date remained, and E14.5 CA1PNs cells were found to display a longer dendritic main branch within the stratum radiatum than both E12.5- and E16.5CA1PNs (E12.5 vs E14.5, P: 0.0056; E14.5 vs E16.5, P: 0.0004, Fig. 6C). Thus, consistent with our previous findings in CA3 (***Marissal et al., 2012***), the dendritic morphology appears to be determined by the temporal embryonic origin, at least in its basic features. Again, this correlation with birthdate is not linear since E12.5 CA1PNs were found to be more similar to E16.5PNs than E14.5CA1PNs, their closer peers in age.

**Figure 6.**
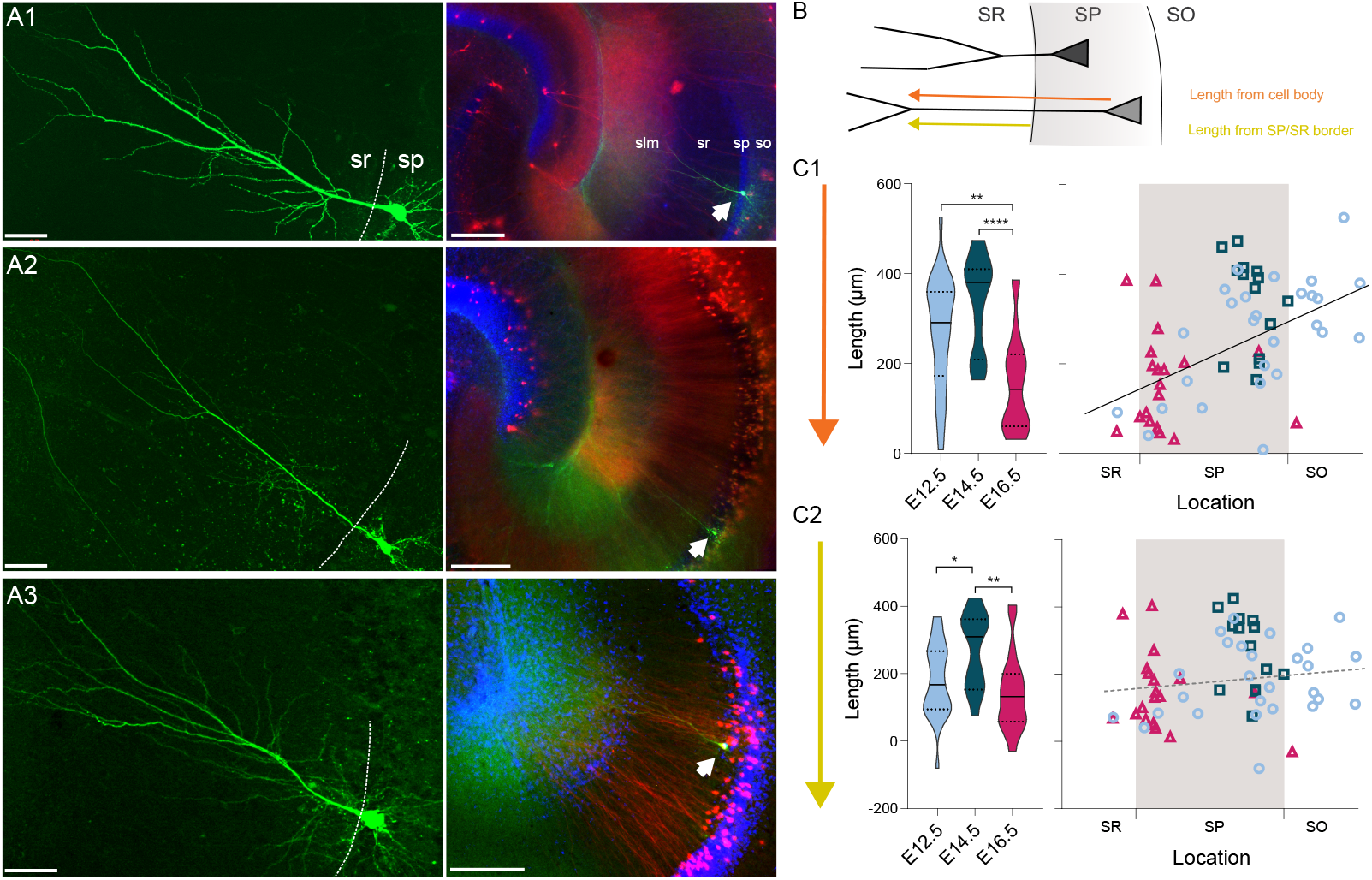
Embryonic origin is a determinant of dendritic morphology. **(A1-3)** On the left, representative examples of CA1 PNs dendritic arborization in neurobiotin-filled cells. The dashed line is drawn at the border between stratum pyramidale (SP) and stratum radiatum (SR). On the right, the same cell as on the left is shown at lower magnification and indicated by a white arrow. Red and blue channels represent Tdtomato and DAPI, respectively. **(A1)** E12.5 PN, **(A2)** E14.5 PN, **(A3)** E16.5 PN. Note that the main dendritic branch bifurcates more distally in E14.5 cells. Scalebars: left, 50 µm; right, 200 µm. so: stratum oriens; sp: stratum pyramidale; sr: stratum radiatum; slm: stratum lacunosum-moleculare. **(B)** Scheme illustrating the primary dendrite length measured from soma or from the border between SP and SR to the first major dendritic bifurcation. SO: stratum oriens. **(C1)** Primary dendrite length from soma to bifurcation in fate mapped CA1PNs. Left, violin plot of the three birthdate groups. Right, scatterplot of the same data against the radial position. E16.5PNs display a reduced dendrite length, in respect to E12.5 (P: 0.0001; CI_95%_ [37.74; 190.36]) and E14.5 PNs (P< 0.0001; CI_95%_ [92.07; 262.82]). The dendrite length from soma is markedly correlated with the soma location (P<0.0001; CI_95%_ [0.318; 0.683]), likely due to the contribution of the dendritic segment within the SP. **(C2)** Primary dendrite length measured between the SP/SR border and the dendrite bifurcation in fate mapped CA1PNs. Left, violin plot of the three birthdate groups. Right, scatterplot of the same data against the radial position. The dendritic length is larger in E14.5 than E12.5 PNs (P_adj_: 0.0056; CI_95%_ [-172.99; -6.14]) and E16 cells (P_adj_: 0.0004; CI_95%_ [36.48; 217.12]). No linear correlation was found with the cell location. Violin present medians (center), interquartile ranges (bounds), minima and maxima. Color-code: E12.5: light blue, E14.5: dark blue, E16.5: magenta. The gray shaded area in scatterplots represents the thickness of the stratum pyramidale. *P < 0.05. **P < 0.01. ****P < 0.0001.

### cFos labelling following the exploration of familiar or novel environments correlates with birthdate

The dorsal CA1 was recently shown to comprise two functional sublayers with deep CA1PNs supporting the formation of landmark-based spatial maps during a novel experience, while superficial CA1PNs were more active in cue-poor conditions and likely to convey self-referenced information (***Danielson et al., 2016; Fattahi et al., 2018; Geiller et al., 2017; Sharif et al., 2020***). We asked whether similar differences could be observed in the ventral CA1 and whether they could reflect developmental origin, given the different connectivity schemes and excitability across CA1PNs with different birthdates. To this aim, we tested the activation of birth dated ventral pyramidal neurons during the exploration of an environment, by the means of cFos immunoreactivity, given the poor accessibility of the ventral hippocampus to imaging and the sparsity of pioneer CA1PNs. cFos is an immediate early gene whose expression does not simply reflect previous activity in labelled neurons but also the induction of activity-related plasticity (***West et al., 2002***).

Ngn2-Cre^ER^-Tdt mice (n=25) induced at E12.5, E14.5 and E16.5 were divided into three groups (Fig. 7A): one explored a cue-enriched arena during 20 minutes for 3 consecutive days, the familiar group (FAM), whereas another group was exposed to the same environment during only one session on the third day, the novel group (NOV). A control group was handled by the experimenter but did not explore any arena, the home cage group (HC). As expected, FAM mice decreasingly explored the arena throughout the 3 consecutive sessions (repeated measures one-way ANOVA, test for trend: P< 0.01, Fig. 7A). Coherently, the distance run by NOV mice was significantly higher than that of the second and third FAM sessions (one-way ANOVA, NOV vs FAM2, P < 0.001; NOV vs FAM3, P < 0.001), but not of the first session (FAM1). As expected, the overall proportion of cFos^+^/DAPI neurons was significantly higher in NOV (10.76 % ± 3.42) and FAM (9.40 % ± 1.31) than the HC condition (5.44% ± 2.45), further confirming that cFos expression in hippocampal neurons is up-regulated upon exploratory activity (Fig. 7B).

**Figure 7.**
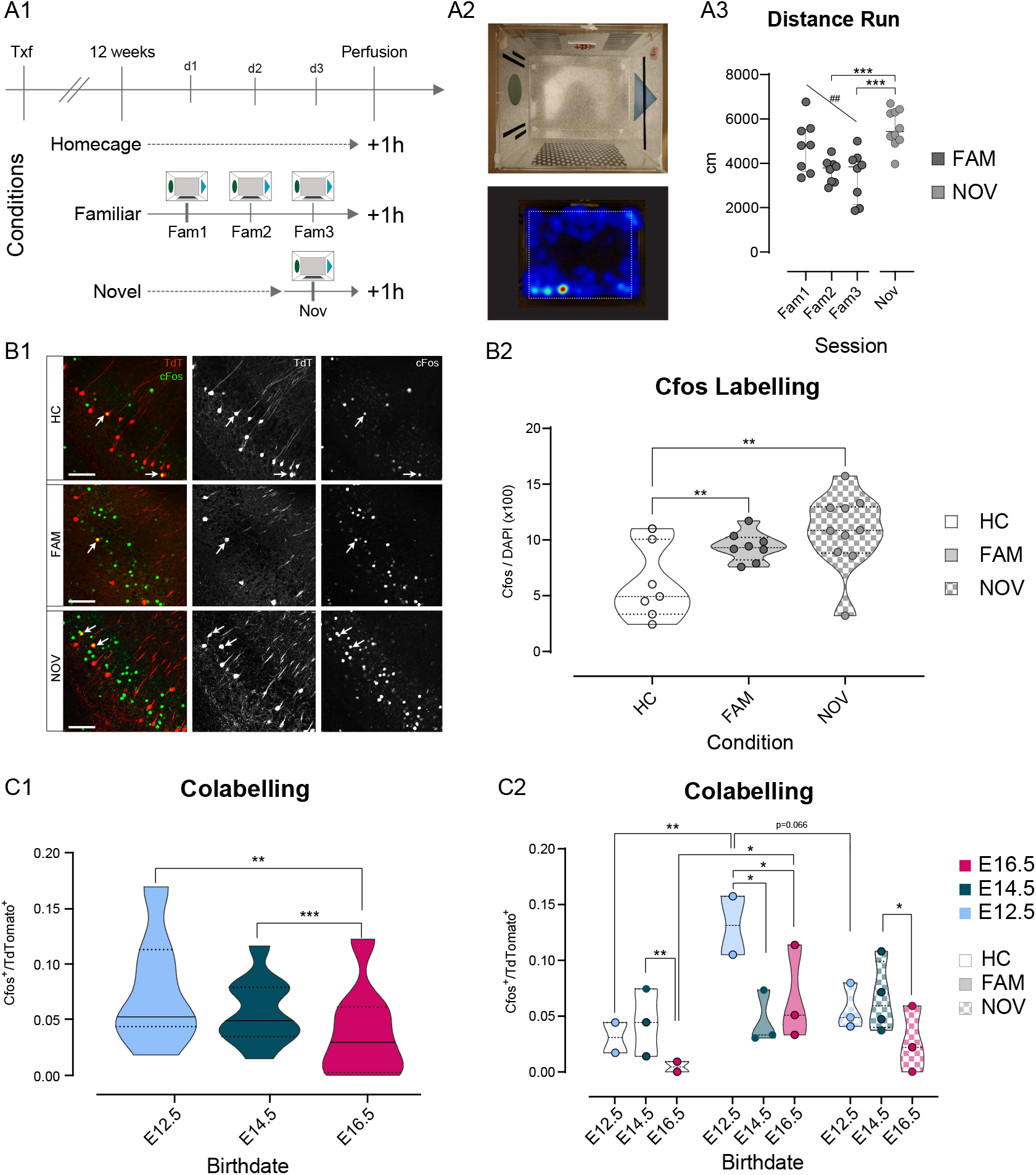
Fate mapped CA1PNs differ in cFos expression following exploratory behavior. **(A1-3)**Experimental paradigm and validation for detecting cFos expression upon exploration of an arena. **(A1)** Schematic representation of the behavioral conditions, with 3 cohorts of tamoxifen-induced Ngn2-Cre^ER^-Ai14 mice: homecage (HC), no exploration, familiar (FAM), repeated exploration (20’) on three consecutive days, and novel (NOV), only one exploration (20’). **(A2)** Top, view from above of the exploration box containing visual, tactile, and olfactory (butanal) cues. **(A3)** In addition, a white noise was played in the experimental room. Bottom, representative occupancy heat map. Quantification of the distance run by animals during exploration of the arena. Upon repeated exposure (Fam1-3), mice explored progressively less the arena (P: 0.0044, CI_95%_ of slope [-242.95;-1077.2]), suggesting that novelty decreases over sessions. Also, NOV did not differ significantly from Fam1 (P_adj_: 0.26), where mice explored for the first time, and was higher than Fam3 (P_adj_: 0.0006, CI_95%_ [973.4; 3264.8]), corresponding to the last exploration. **(B1-2)** The expression of cFos in ventral CA1 is higher upon exploration of a novel environment, than after repeated exposures. **(B1)** Immunohistochemistry anti-cFos, performed on three tamoxifen-induced mice (E14). Top, HC; middle, FAM; bottom, NOV condition. Each shows the merged Tdt, cFos and DAPI image (left), Tdt (center), cFos (right). Note the increase in cFos signal from HC to FAM and NOV. Insert: magnification on two cells, one strongly cFos+, the other co-expressing Tdt and cFos. Scalebar: 100 µm. **(B2)** Fraction of cFos^+^ cells per animal. In both FAM (P_adj_: 0.0006, CI_95%_ [-5.78; -1.99]) and NOV (P_adj_: 0.0006; CI_95%_ [-7.86; -2.54]) animals, cFos immunoreactivity is increased compared to HC. **(C1-2)** Quantification of the co-expression of cFos and Tdt in fate-mapped CA1PNs. **(C1)** Fraction of cFos^+^/Tdt^+^ cells by birth date group, regardless of the condition. E16 cells display fewer co-labelled cells than E12 (P_adj_: 0.0022, CI_95%_ [0.0097; 0.0483]) and E14 (P_adj_: 0.0009, CI_95%_ [0.0137; 0.0459]). **(C2)** Fraction of cFos^+^/Tdt^+^ cells by birth date and per condition. Note that within the FAM condition, cFos^+^/Tdt^+^ cells are more abundant in E12-FAM than E14-FAM (P_adj_: 0.008, CI_95%_ [0.0284; 0.135]) and E16-FAM (P_adj_: 0.0325, CI_95%_ [0.0236;0.136]). Within E12 birth date group as well, E12-FAM fraction of co-labelled cells is higher than E12-HC (P_adj_< 0.0001, CI_95%_ [-0.151; -0.049]), and roughly higher than E12-NOV (P_adj_< 0.066, CI_95%_ [0.0149; 0.123]). In addition, E14-NOV is greater than E16-NOV (P_adj_< 0.015, CI_95%_ [0.0179; 0.0797]). Violin present medians (center), interquartile ranges (bounds), minima and maxima. Color-code: E12: light blue, E14: dark blue, E16: magenta. *P < 0.05. ##, **P < 0.01. ***P < 0.001.

To study how cFos activation may vary according to a neuron’s embryonic origin, we first computed the proportion of cFos^+^/Tdt^+^ cells by birth date group, regardless of the condition (Fig. 7C). Doing so, we observed that E16.5 mice displayed significantly lower proportions of co-labelled cells than both E12.5 (95% Confidence interval or CI95 [0.0097; 0.0483], P < 0.01) and E14.5 (CI95 [0.0137; 0.0459], P < 0.001). Hence, late-born E16.5CA1PNs are less prone to express cFos, as expected from the previously reported global difference in overall recruitment between deep and superficial CA1 PNs (***Mizuseki et al., 2011***). Using the same approach as for retrograde tracing analysis, we examined the cFos^+^/Tdt^+^ fraction per condition per group. We found that pioneer E12.5CA1PNs were highly likely to express cFos following habituation (E12.5-FAM), as compared across conditions within the same birth date group (E12.5-FAM vs E12.5-HC, P < 0.001; E12.5-FAM vs E12.5-NOV, P: 0.066) and across birthdates for the same FAM condition (E12.5-FAM vs E14.5FAM, P < 0.01; E12.5-FAM vs E16.5-FAM, P < 0.05, Fig. 7C). Although more than twice less than for E12.5CA1PNs, E16.5CA1PNs were also significantly more likely to express cFos in FAM than HC conditions. These cells were overall very unlikely to express cFos, especially in the HC condition where co-labelling was significantly lower than observed in E14.5 mice (E16.5-HC vs E14.5-HC, P < 0.01). Importantly, in the NOV condition, earlier-generated cells showed more cFos^+^ activation than later born ones, though this only appeared significant for E14.5 CA1PNs (E14.5-NOV vs E16.5-NOV, P < 0.05; E14.5-NOV vs E12.5-NOV, P: 0.165).

Taken together, these results are in line with previous research, in that deep CA1PNs are more likely to express IEGs than superficial ones upon exposition to a novel environment (***Kitanishi et al., 2009***). Yet, pioneer E12.5CA1PNs form a distinct subpopulation more specifically expressing cFos after familiarization to the environment. Hence, these last results indicate a different involvement in novelty detection of pyramidal neurons in the ventral CA1 according to their developmental origin.

## Discussion

In this study, we have employed a manifold description of CA1PNs with different birthdates in the adult hippocampus to study the origin of diversity at single-cell level. We have found that the now well-established subdivision of the CA1 pyramidal layer into two distinct sublayers, a deep and superficial one, masks a much greater heterogeneity, which logic can be exposed when considering the temporal origin of individual neurons. This heterogeneity encompasses several morphophysiological properties and translates in the propensity of cells to be recruited in familiar or novel environments. Furthermore, we find that pioneer CA1PNs generated at the earliest stages of glutamatergic cell neurogenesis are prone to be recruited in familiar environments, forming a subpopulation with distinct anatomical distribution, morphophysiological features and wiring.

### Birthdate rather than birth order is a better predictor of diversity

Several studies converge in suggesting that the CA1 pyramidal layer comprises two subtypes of CA1PNs based on their radial positioning. We would like to propose that the diversity of CA1PNs is even better predicted by birthdate than position. More specifically, we found that the following metrics were better reflecting birthdate than radial positioning based on the lack of correlation with soma position and the existence of a significant correlation with birthdate: (1) the length of primary apical dendrite in the stratum radiatum; (2) the input resistance, sag current and firing rate in response to depolarizing current injections; (3) the E/I ratio; (4) the short-term facilitation of evoked synaptic inhibitory currents. The two main metrics that linearly correlate with layering reflect the local microcircuit integration of CA1PNs as they are the frequency of sEPSCs and the evoked IPSC decay.

It was previously reported that both ectopic CA1PNs with a soma located in the *stratum oriens* received a higher rate of sEPSCs than cells located within the *stratum pyramidale* (***Cossart et al., 2000***). Similarly, we find that CA1PNs located closer to the *stratum oriens* receive sEPSCs at a higher rate. This could reveal either a slicing artefact or the contribution of glutamatergic inputs from the CA1 axonal collaterals arborizing in that area. The faster kinetics of evoked IPSCs may reflect the increased contribution of parvalbumin basket cell inputs to evoked IPSCs as cells move towards the oriens. Indeed, the alpha1-containing postsynaptic GABAA-Rs contributing to PV synapses were shown to display faster kinetics than those formed by CCK basket cells (***Thomson et al., 2000***). In light with studies indicating specified microcircuitry among deep versus superficial principal cells and PV basket cells, we have found that the PV staining was denser towards the *stratum oriens*. However, within the deep CA1PNs, cells born at E12.5 were exceptionally less decorated with putative PV contacts than deep CA1PNs born 2 days later. Overall, we observed that E12.5 cells were more similar to E16.5 CA1PNs regarding their lower intrinsic excitability, dendritic morphology, lower PV coverage and higher E/I ratio. As E16.5CA1PNs, they can even be found in the superficial layer and even in the stratum radiatum. Altogether, one could foresee that the main source of activation for both E12.5 and E16.5 CA1PNs would be through synaptic glutamatergic excitation. However, it is likely that the main glutamatergic inputs onto E16.5 and E12.5 CA1PNs differ. While the main excitatory drive onto E16.5CA1PNs, given their location likely originates in CA3, future studies are needed to determine whether E12.5 cells receive specific inputs. Such studies remain at the moment technically challenging due to the poor accessibility of E12.5 cells to genetic manipulation using our fate-mapping strategy.

These cells also differ in their extrahippocampal outputs. Accordingly, these cells could also be discriminated based on their cFos expression following spatial exploration with E12.5, like E14.5 CA1PNs being more likely to express cFos during random exploration than E16.5CA1PNs. Future studies are needed to determine the conditions favoring cFos expression in E16.5CA1PNs. These may require analyzing these cells in conditions of elevated anxiety, social interactions or goal-directed behaviors (***Ciocchi et al., 2015; Jimenez et al., 2018; Okuyama et al., 2016***). If E12.5CA1PNs and E14.5CA1PNs displayed comparable cFos expression levels, as expected from previous work (***Mizuseki et al., 2011***), the former signals familiarity and the latter novelty, suggesting that the mechanisms supporting hippocampal representation of novelty may require cells with a higher intrinsic neuronal excitability (***Epsztein et al., 2011***).

If our results are mostly in agreement with previous reports, we were not able to observe any significant correlation between laminar position and sag current amplitude, resting membrane potential or input resistance in our dataset. Only E12.5CA1PNs could be distinguished by their lower sag amplitude value and input resistance. This apparent discrepancy may have several explanations. First, our experiments were not performed in the presence of blockers for other postsynaptic membrane currents and of antagonists for all metabotropic and ionotropic GABA and glutamate receptors (***Jarsky et al., 2008; Lee et al., 2014; Maroso et al., 2016; Masurkar et al., 2017***). Second, we focused on ventral and not dorsal hippocampus (***Maroso et al., 2016***), and cells were sampled evenly across the transverse axis, while radial correlations could only be revealed at fixed positions along the proximo-distal axis (***Masurkar et al., 2017***). We were also surprised not to observe any bursting cell (***Jarsky et al., 2008; Graves et al., 2012***), but again these are mainly present closer to the subiculum and were reported in slices from juvenile (P15-17 (***Jarsky et al., 2008***), P25-28 (***Graves et al., 2012***)) rats using a gluconate-containing intracellular solution (***Kaczorowski et al., 2007***). Last, while we could observe a linear correlation between soma location and dendritic morphology as reported previously (***Graves et al., 2012; Jarsky et al., 2008; Li et al., 2017; Lee et al., 2014***), this was no longer observed when considering only the dendritic portion within the stratum radiatum. When computing this metric, E14.5CA1PNs could be distinguished from E12.5 and E16.5 CA1PNs by their longer primary apical segment. Also, one needs to consider one major limitation of the fate mapping approach employed here when interpreting the results of the present study. Indeed, our labelling is based on the Cre-dependent and tamoxifen-induced expression of a fluorescent reported protein in Ngn2 expressing progenitors. This is a stochastic process, and a variable and sparse subset of progenitors get labelled. We are only describing small numbers and the most distinctive features are more likely to show up as significantly different whereas other metrics may require denser sampling.

Altogether, our results globally indicate that the order of neurogenesis does not imprint a continuous gradient in the specification of cell laminar positioning and diversity. Instead, these are predetermined by the specific temporal origin of individual cells. Interestingly, the notion that early specification has a stronger importance than final layering in the determination of CA1PNs intrinsic properties is supported by recently observations with a transgenic mouse model of lissencephaly where CA1 lamination is impaired while cell identity is relatively spared (***D’Amour et al., 2020***). Therefore, the recently uncovered system of parallel information processing in CA1, with two information streams segregated through distinct inhibitory domains (***Soltesz and Losonczy, 2018***), may need to consider this additional level of complexity.

The mechanisms by which diversity among CA1PNs may be temporally-regulated during neurogenesis remain to be determined. Increasing evidence indicates that glutamatergic neurons are issued from the same multipotent progenitors and that fate distinctions are mostly temporally controlled (***Jabaudon, 2017***) (but see Franco et al. (***2012***)). Interestingly, progenitor potential was shown to be progressively, temporally restricted, with early cortical progenitors being multipotent in comparison to later ones (***Lodato et al., 2015***). In combination with genetic predetermination, single-cells display tightly orchestrated sequences of spontaneous activity patterns (***Allene et al., 2012***), which in turn may contribute to the maturation of physiological specificity. Therefore, cells with similar birthdates will follow similar activity development schedules. Interestingly, both Ih and Rin, two parameters that segregate between CA1PNs with different birthdates, are developmentally regulated, both progressively decreasing as cells mature (***Dougherty, 2019***). As such, they are ideal proxys of neuronal maturation stage. Given that the earlier born CA1PNs can be distinguished by lower values of Ih and Rin, one could propose that these cells could maintain into adulthood the advance in maturation originating from their early birth date.

### Pioneer cells are a different population, possible roles?

This study shows that neurons born at the earliest phases of CA1PNs neurogenesis display distinct somatic distribution, connectivity, morpho-physiological properties as well cfos expression patterns. This distinguishes them from slightly later born neurons, the E14.5CA1PNs that sit mainly in the deep sublayer of the *stratum pyramidale*. We argue that these cells form a distinct subpopulation. The first striking property of E12.5CA1PNs is their scattered distribution. They can be found at ectopic positions in the *stratum oriens* as well as, more rarely, within the stratum radiatum, suggesting that these cells may also comprise the previously described radiatum “giant cells”, with privileged CA2 input and output to the olfactory bulb (***Gulyás et al., 1998; Nasrallah et al., 2019***). In fact, cumulative distribution plots indicate that roughly half of the E12CA1PNs in the ventral (and dorsal) hippocampus are found within less than 100 µm distance from the stratum radiatum while the other half distributes closer to *stratum oriens* more than 100 µm away from that border. It is unlikely that such observation results from unspecific labeling from our method. First, the population of E12.5CA1PNs labeled using this method shares many characteristics despite this layer dispersion. Second, this unique arrangement was also recently observed for early born granule cells (***Save et al., 2019***) and dispersion of early generated glutamatergic cell cohorts was previously suggested using other methods including autoradiographic (***Caviness, 1973***) and retrovirus labeling (***Mathews et al., 2010***). Somehow similarly, even if preferentially located in the deep stratum pyramidale, early born CA3 pyramidal cells could be found anywhere from *stratum oriens* to *lucidum* (***Marissal et al., 2012***). It is therefore possible that pioneer cohorts of PNs would display a different migration mode, resulting from the low cellular density when entering the hippocampus in addition to other more specific mechanisms, like an absence of radial glia bending (***Xu et al., 2014***), that remain to be specifically studied. Regardless, despite this almost even positioning, we observed that E12.5CA1PNs displayed specific input connectivity schemes, including a lower number of putative PV contacts associated with a higher E/I ratio and a facilitation of the evoked IPSC amplitude. We hypothesize that these three measurements may reveal the same feature, namely the lower somatic inhibitory coverage of E12.5CA1PNs. Indeed, facilitating inhibitory responses were found to originate from dendrite-targeting interneurons while perisomatic cells would tend to generate transient inhibitory inputs (***Pouille and Scanziani, 2004***). This lower PV innervation is comparable with E16.5CA1PNs and contrasts with their early-born peers, the E14.5CA1PNs, which display higher perisomatic PV staining. Whether these CA1PNs are contacted by specific glutamatergic inputs stays an open question that remains an experimental challenge with our fate-mapping strategy. According to the temporal matching rule observed in the hippocampus (***Deguchi et al., 2011***), and the fact that some of them are found in the stratum radiatum and oriens, one may expect these cells to receive preferential innervation from CA2, the earliest generated portion of the hippocampal circuit (***Caviness, 1973***). The significantly higher E/I ratio received by these cells suggests that they may be preferentially recruited by synaptic excitation, since they are otherwise less intrinsically excitable (lower Rm, low firing rate in response to long depolarizing current injections, lower sag). Interestingly, the same lower excitability (across similar metrics), combined with lower levels of PV inputs was observed in early born CA1 GABAergic neurons and DG granule cells (***Bocchio et al., 2020; Save et al., 2019; Gupta et al., 2012***), again indicating similar fates in the adult across different subtypes of pioneer neurons.

Ventral E12.5CA1PNs are also remarkable in terms of output since they globally sent more projections to the target regions studied here and did not display any preferred projection pattern in contrast to their later born peers. Of note they were significantly more likely to target the Amy and mPFC than both E14.5 and E16.5 cells. These results would suggest that E12.5CA1PNs may form a sparse population with multiple projections. Triple projection ventral CA1 cells targeting the mPFC, Amy and NAcc were shown to be highly active and in particular during SWRs (***Ciocchi et al., 2015***). Unfortunately, the sparsity of our labeling method currently prevents opto-tagging E12.5CA1PNs as well as testing the hypothesis that they may be comprised of multiple projection neurons.

## Conclusion

We have uncovered a novel population of pioneer cells adding to the diversity of CA1 glutamatergic neurons. Based on their specific properties, we would like to propose a general role for ventral pioneer CA1PNs in the consolidation or retrieval of recent experience. Future work examining their recruitment in SWRs (***Ciocchi et al., 2015***), their possible CA2 inputs (***Kohara et al., 2014; Nasrallah et al., 2019; Valero et al., 2015***) and their likely multiple projection targets will certainly inform about this possibility. In any case, the present results indicate that the radial subdivision of the CA1 pyramidal layer into two functional sublayers needs to be revisited by considering the time of neurogenesis. The strongest reliance of CA1 principal neuron diversity on temporal origin or initial genetic blueprint than position may render these cells more resilient to diseases resulting in heterotopias and mislayering, as recently reported (***D’Amour et al., 2020***).

## Methods and Materials

### Animals

All protocols were performed under the guidelines of the French National Ethics Committee for Sciences and Health report on “Ethical Principles for Animal Experimentation” in agreement with the European Community Directive 86/609/EEC under agreement #01 413.03. All efforts were made to minimize pain and suffering and to reduce the number of animals used. Animals were maintained with unrestricted access to food and water on a 12-hour light cycle, and experiments were conducted during the light portion of the day. To mark glutamatergic neurons generated during different times of embryogenesis, we use the technique of inducible genetic fate mapping, see (***Marissal et al., 2012; Save et al., 2019***). In brief, double heterozygous *Ngn2-CreER*^*™/-*^*/Ai14-LoxP*^*+/-*^ mice (referred as Ngn2-Cre^ER^-Ai14 in the text for simplicity) were obtained by crossing *Ngn2-CreER*^*™/-*^*/Ai14:LoxP*^*+/+*^ male mice with 7-8-week old wild-type Swiss females (C.E Janvier, France). To induce Cre activity during embryogenesis, tamoxifen was delivered to pregnant mothers (Sigma, St. Louis, MO; 2 mg per 30 g of body weight of tamoxifen solution, prepared at a concentration of 10 or 20 mg/mL in corn oil) at embryonic days E12.5, 14.5, 16.5.

### Slice preparation for *ex vivo* electrophysiology

Ngn2-Cre^ER^-Ai14 mice of either sex aged between post-natal week 4 (W4) and W11 treated with tamoxifen at E12.5, E14.5 or E16.5 were used for experiments. First, they underwent deep anesthesia by intraperitoneal xylazine/ketamine injection (Imalgene 100 mg/kg, Rompun 10 mg/kg) prior to decapitation. 300 µm-thick horizontal slices were cut with a Leica VT1200 Vibratome using the Vibrocheck module in ice-cold oxygenated modified artificial cerebrospinal fluid with the following composition (in mM): 126 CholineCl, 26 NaHCO3, 7 MgSO4, 5 CaCl2, 5 D-glucose, 2.5 KCl, 1.25 NaH2PO4). Slices were then transferred for rest at room temperature (at least 1 h) in oxygenated normal aCSF containing (in mM): 126 NaCl, 3.5 KCl, 1.2 NaH2PO4, 26 NaHCO3, 1.3 MgCl2, 2.0 CaCl2, and 10 D-glucose, for a total of 300 ± 10 mOsm (pH 7.4). A total of 72 cells were successfully recorded from 34 mice (n_E12.5_ = 22, n_E14.5_ = 27, n_E16.5_ = 23). 63 cells were used for morphological analysis.

### *Ex vivo* patch clamp recordings

Patch clamp recordings in adult slices were performed using a SliceScope Pro 1000 rig (Scientifica) equipped with a CCD camera (Hamamatsu Orca-05G) and with a X-Cite 120Q (Excelitas Technologies Corp.) fluorescence lamp. Recordings were performed from visually identified Tdt+ cells. Patch electrodes (4-7 MΩ resistance) were pulled using a PC-10 puller (Narishige) from borosilicate glass capillaries (GC150F10, Harvard Apparatus). Slices were transferred to a submerged recording chamber and continuously perfused with oxygenated ACSF (3 mL/min) at 32 °C. Electrophysiological signals were amplified, low-pass filtered at 2.9 kHz and digitized at 10 kHz with an EPC10 amplifier (HEKA Electronik) and acquired using the dedicated software PatchMaster. Neurons were kept at -65 mV throughout the session and recordings started 5-10 minutes after access. For current clamp experiments, the following intracellular solution was used (in mM): 130 K-MeSO4, 5 KCl, 5 NaCl, 10 HEPES, 2.5 Mg-ATP, 0.3 GTP, and 0.5% neurobiotin (∼280 mOsm, 7.3 pH). Capacitance compensation and bridge balance were performed before recording and adjusted periodically. Liquid junction potential was not corrected. The resting membrane potential was not measured. For voltage clamp experiments, the solution consisted of (in mM): 130 CsGluconate, 10, HEPES, 5 CsCl, 5 Na2Phosphocreatine, 2 MgATP, 1 EGTA, 0.3 Na3GTP, 0.1 CaCl2 and 5% neurobiotin ∼290 mOsm, 7.3 pH). Uncompensated series resistance was calculated postacquisition and considered acceptable if below 30 MΩ and if variations were lower than 20%. Liquid junction potential correction (−13 mV) was applied. Excitatory and inhibitory postsynaptic currents were recorded at -86 mV (EGABA) and 0 mV (EGlu) respectively, over 9 sweeps of 20s. A -5 mV step (5 ms duration) was used to monitor series resistance at the beginning of each sweep. For Schaffer collateral stimulation, a bipolar electrode made by two twisted nichrome wires and connected to a DS2A Isolated Voltage Stimulator (Digitimer) was used to deliver 0.2 ms-long stimuli in the stratum radiatum of CA1 (between the recorded cell and CA3). Stimulation intensity was set to 2X the minimum intensity capable of eliciting a postsynaptic response (tested between -86 mV and -78 mV). Paired pulse ratio was assessed by two stimulations separated by 50 ms.

### Analysis of *ex vivo* patch clamp recordings

Analysis of intrinsic membrane properties was performed using custom-made MATLAB scripts. Traces were filtered using the sgolay MatLab function, unless firing properties were being analysed. The input membrane (R_m_) resistance was calculated from the slope of steady-state voltage responses to a series of subthreshold current injections lasting 500 ms (from -100 pA to last sweep with a subthreshold response, 10 or 20 pA step size). The membrane time constant (τ_m_) was estimated from a bi-exponential fit of the voltage response to a small (−20 pA) hyperpolarizing pulse. Capacitance (C_m_) was calculated as τ_m_/R_m_. The sag potential was calculated by injecting a 1 s-long negative current step (−200 pA) as follows: Sag_-200pA_=Δ*V* _steady-state_-Δ*V* _peak_ (subtraction of the steady-state potential at halfresponse (averaged over 45 ms) to the minimum peak potential at the beginning of the response). The rebound potential was computed, in absence of rebound action potentials, by subtracting the depolarization peak after the end of the hyperpolarization to the baseline potential: Rebound_-200pA_=Δ*V* _peak_-Δ*V* _baseline_. Firing curves were determined by applying an increasing range of 1-s long depolarizing current injections (up to 500 pA, +40 pA step). The rheobase (in pA) was defined as the minimum current value necessary to elicit an action potential. The first spike in response to a positive current injection was used to determine: the threshold potential (the peak of the second derivative), the action potential amplitude and fast afterhyperpolarization (both calculated from threshold potential), and the half-width (width at half-amplitude between the threshold potential and the peak of the action potential). Analysis of spontaneous postsynaptic currents (sPSCs) was performed using MiniAnalysis (Synaptosoft). Traces were filtered with a LoPass Butterworth (cutoff frequency 1000 Hz) and PSC amplitude and frequency were calculated over 2 minutes (total of 6 sweeps, 20s). The mean PSC event was analysed with a custom-made MATLAB script to compute rise time (10-90% of the ascending phase); decay time (90-37% of the descending phase); decay τ (from a monoexponential curve adjusted to best fit the trace); half-width (width at half-amplitude between PSC peak and baseline); area (using trapz MATLAB function). Mean stimulation-evoked PSC amplitude, area and kinetics were calculated as detailed above. Paired pulse ratio was measured by calculating the amplitude of two 0.2 ms-long stimuli at 50 Hz as follow: PPR=Peak_2_/Peak_1_. The ratio between excitatory and inhibitory PSC (E/I ratio) was calculated by dividing frequency and amplitude values.

### Morphological analysis of neurobiotin-filled cells

Slices containing neurobiotin-filled cells were fixed overnight at 4 °C in PFA (4%), rinsed in PBS containing 0.3% Triton X-100 (PBS-T) and incubated overnight at room temperature in streptavidin-AlexaFluor488 (1:1000 in PBS-T). Slices were mounted using Vectashield mount medium containing DAPI (VectorLabs). Post-hoc analysis was performed using an AxioImager Z2 microscope equipped with Apotome module (Zeiss). The co-localisation of neurobiotin and TdTomato was verified systematically for every recorded cell. Neurobiotin-filled neurons were selected when the apical dendrites were clearly visible and uncut. Stacks of optical sections were collected for these cells. The primary dendrite was identified as the portion of between the soma and a bifurcation generating secondary dendrites of roughly equal size. The length of the primary dendrite was measured by approximating its shape to 1-3 linear segments.

### Quantification of cell location in stratum pyramidale

For patch clamp recordings, the cell location was measured from the border between the *stratum pyramidale* (SP) and the *stratum radiatum* (SR) and normalized to the thickness of the SP, meaning that values close to 0 correspond to the proximity of SP/SR border and values close to 1 to *stratum oriens* (SO)/SP border. For the overall quantification of Ngn2 cell location, the cell location was measured from the border between the SP and the SR and expressed in micrometers.

### Histological processing and immunohistochemistry

After deep anesthesia with a ketamine (250 mg/kg) and xylazine (25 mg/kg) solution (i.p.), animals were transcardially perfused (1 mL/g) with 4% paraformaldehyde in saline phosphate buffer (PBS). Brains were post-fixed overnight, then washed in PBS. For Ctb tracing, a VT1200 Vibratome (Leica) was used to cut coronal slices. Slice thickness was 100 µm for the injection site and 70 µm for the hippocampus. In a subset of experiments, CTB-AlexaFluor647 signal was amplified by immunohistochemistry. Sections were incubated overnight with primary Cholera Toxin beta polyclonal antibody (rabbit; 1:1000, RRID: AB_779810, Invitrogen) diluted in PBS-T and for approximately 1h30 with donkey antirabbit secondary antibody AlexaFluor647 (1:500, Jackson Immunoresearch, 711-606-152) in PBS-T. For cfos immunostaining, a similar procedure was employed. Coronal slices (50 µm thickness) obtained with a HM450 sliding Microtome (Thermo Scientific, Waltham, MA) were rinsed 3 times in PBS containing 0.3% Triton X (PBS-T) and incubated overnight at 4° C with a solution containing 5% goat serum (GS) and anti-cFos (rabbit; 1:5000, AB_190289, Abcam) diluted in PBST. The following day slices were exposed to a goat anti-rabbit AlexaFluor647 secondary antibody (1:500, Jackson Immunoresearch, 111-606-144) in PBS-T. Post-hoc analysis was performed from image stacks (1.5 µm interval, 7 images) obtained using a Zeiss LSM-800 system. Slices were mounted using Vectashield mount medium containing DAPI (VectorLabs). Post-hoc analysis was performed from image stacks obtained using a Zeiss LSM-800 system equipped with emission spectral detection and a tunable laser providing excitation range from 470 to 670 nm.

### Quantification of putative PV-expressing synaptic boutons

For parvalbumin (PV) putative puncta detection, we employed a similar immunostaining procedure as described above. Goat anti-PV primary antibody (1:1000, Swant, pvg-214, AB10000345) and donkey anti-goat AlexaFluor488 secondary antibody (1:500, Jackson Immunoresearch, AB_2336933) were used. Stacks (0.06 x 0.06 x 0.410 µm) centered on the soma of TdTomato+ cells (E12.5: 88 cells; E14.5: 106 cells; E16.5: 44 cells) were acquired with a Zeiss LSM-800 microscope using a Plan-Apo 40x/1.4 oil objective. Volume overlaps between PV+ boutons and Tdt+ somata were calculated using a custom-made MatLab script. Values were normalized by the number of Z-steps to control for possible differences in soma size or experimental variability.

### Stereotaxic injections for retrograde tracing

Ngn2-Cre^ER^-Ai14 adult mice (age > P50) of either sex were anaesthetized using 1-3% isoflurane in oxygen. Analgesia was provided with buprenorphine (Buprecare, 0.1 mg/kg). Lidocaine was applied by cream or subcutaneous injection before the incision for additional local analgesia. Mice were fixed to a stereotaxic frame with a digital display console (Kopf, Model 940). Under aseptic conditions, an incision was made in the scalp, the skull was exposed, and a small craniotomy was drilled over the target brain region. A 200-400 nl volume of 0.1% AlexaFluor647-conjugate Cholera Toxin subunit b (CTB, Thermofisher Scientific) was delivered using a glass pipette pulled from borosilicate glass (3.5” 3-000-203-G/X, Drummond Scientific) and connected to a Nanoject III system (Drummond Scientific). The tip of the pipette was broken to achieve an opening with an internal diameter of 30-40 µm. Stereotaxic coordinates were based on a mouse brain atlas (Paxinos and Franklin, 3rd edition). All coordinates are indicated in Table 1 in millimeters, and examples of the histological recovery of the injection sites are displayed in Fig. 4-Supplement1. Antero-posterior (AP) coordinates are relative to bregma; medio-lateral (ML) coordinates are relative to the sagittal suture; dorso-ventral (DV) coordinates are measured from the brain surface. Mice were perfused 12-15 days later, to allow sufficient CTB uptake and transport from the synaptic terminals.

**Table 1.** Stereotaxic coordinates of CA1PNs target regions.

| Target Region | AP | ML | DV |
| --- | --- | --- | --- |
| **NAcc** | +1.8/+2.0 | -0.6/-0.45 | -4.0 |
| **Amy** | -1.4 | -3.3 | -4.3 |
| **mPFC** | +2.0 | -0.35 | -1.8 |
| **LHA** | 1.34/-1.45 | -0.65/-1.0 | -4.8/-4.0 |
| **LS** | +0.6/0.7 | -0.4/-0.35 | -2.6/2.55 |

Abbreviations: NAcc, Nucleus Accumbens (shell), Amy: Amygdala; mPFC, medial prefrontal cortex; LHA, lateral hypothalamic area, LS: lateral septum.

### Analysis of retrograde tracing

To confirm that injections were successful, each injection site was visually inspected and compared to the Paxinos mouse brain atlas. The occurrence of co-labelling with Ctb among Tdt^+^ cells was verified and the ratio of Ctb^+^/TdT^+^ cells calculated manually. This resulted in the generation for each injected animal of a binary vector of length L equal to the total count of Tdt^+^ cells and as many “1” entries as the number of identified Ctb^+^/TdT^+^ cells. Then, vectors corresponding to mice in the same birthdate group and same target region were concatenated. Finally, a bootstrap resampling approach was used to compute the pairwise comparisons (total number of tests = 45). In brief, each of two vectors A and B were randomly resampled with replacement for 10000 times. At each iteration, the difference between the two vector means (Δ_µ_= µ_A_-µ_B_) was calculated and stored. At the end of the resampling, the 99.9% confidence interval (CI) of the distribution of (Δ_µ_) was computed. The difference between vectors A and B was considered significant if the CI did not include the value 0.

### cFos expression upon exploration

A total of 25 tamoxifen-treated Ngn2-Cre^ER^-Ai14 mice between 10-12 weeks old of either sex were used. Prior to the experiment, animals were single-housed and divided in 3 groups, named homecage (HC), familiar (FAM) and novel (NOV), see Table 2. Group HC was carried to the experimental room, handled by the experimenter during 5-10 minutes for three consecutive days (D1, D2, D3) and perfused 1h after handling on D3. Group FAM was exposed to an exploration chamber for 20 minutes from D1 to D3, and returned to their home cage immediately after. Group NOV was handled during D1 and D2 and left in the exploration chamber for one single 20-minute session on D3. FAM and NOV were both perfused 1h after exploration. The chamber consisted of a transparent plastic rat cage sized 435×290mm containing visual, tactile, and olfactory (butanal) cues. A white noise (20/30 dB) was played in the experimental room for the duration of the exploration and low lighting (∼25 lux) was centered over the box. Mice position during each session was recorded with a Basler Ace camera (Basler AG, Ahrensburg) and tracked with EthoVision XT 11 (Noldus, Leesburg, VA) software.

**Table 2.** Summary of adult Ngn2-Cre^ER^-AI14 mice used in post-exploration cFos analysis.

| N° animals | HC | FAM | NOV | Total |
| --- | --- | --- | --- | --- |
| E12.5 | 2 | 2 | 3 | **7** |
| E14.5 | 3 | 3 | 4 | **10** |
| E16.5 | 2 | 3 | 3 | **8** |
| Total | 7 | 8 | 10 | **25** |

### Analysis of cFos expression upon exploration

Tdtomato^+^ somata were segmented with Fiji Trainable WEKA Segmentation plugin. Two custommade Cellprofiler pipelines were used on DAPI and AlexaFluor647 (cFos) channels (***Carpenter et al., 2006***). (i) To provide an estimation of DAPI cell density, a single 2D image (from the middle of the stack) was segmented using a Minimum Cross Entropy threshold. (ii) Segmentation of cFos^+^ nuclei was first achieved by applying the Robust Background thresholding method on each single 2D image composing the stack and converting them into binary masks. Then, these masks were restacked and somas were identified with the 3D Object Counter Fiji plugin across the depth of the stack. To determine TdTomato and cFos colocalization, a custum-made MatLab script was used. In a nutshell, matrices representing TdTomato and cFos binary masks were multiplied, generating a new binary image of areas presenting overlap between the two channels. Finally, each of these areas, which represented putative colocalized cells, was manually inspected for confirmation. At the end of this procedure, we applied the same procedure as described for the analysis of retrograde tracing. We computed for each animal a binary vector of length L equal to the total number of segmented Tdt^+^ cells and as many “1” entries as the number of identified cFos^+^/Tdt^+^ cells. Then, vectors corresponding to mice in the same birthdate group and same behavioral condition were concatenated. Finally, a bootstrap resampling approach was used to compute the pairwise comparisons (total number of tests = 18). The 95% confidence interval (CI) of the distribution of the bootstrapped differences of means (Δ_µ_) was computed. The difference between vectors A and B was considered significant if the CI did not include the value 0.

### Statistical Analyses

Statistical analysis was done using Prism (GraphPad) and costum-made MatLab scripts. In patch clamp experiments, outliers were removed (ROUT method, Q = 0.1%). For comparing input-output firing curves, two-way ANOVA was used. When comparing birthdate groups for a given measure, median-based bootstrap resampling was used to compute pairwise comparisons, subsequently corrected with Holm-Bonferroni method to control for family-wise error rate, except for morphological measures where mean and Sidak’s correction were used instead. The correlation with cell location was tested using bootstrap resampling. In the analysis of exploratory behavior, repeated-measures and ordinary one-way ANOVA were used to compare distance run in FAM condition and FAM vs NOV, respectively. The volume overlap of putative PV boutons is tested among birthdates with one-way ANOVA with Tukey’s correction for multiple comparisons.

## Supporting information

Statistics Tests

## Acknowledgments

This work was supported by the European Research Council under the European Union’s Horizon 2020 research and innovation programs (grant agreement no. 646925) as well as by the Agence Nationale pour la Recherche (ANR, Programme Blanc bilatéraux, ANR-13-ISV40002-01 “EbGluNet” and HIPPOPLAST, JTC-2017-021) and by the Fondation Bettencourt Schueller (Prix des Sciences de la Vie). A.B. and R.C. are supported by the Centre National de la Recherche Scientifique (CNRS). We thank Pr. David Anderson for providing the Ngn2-CreER^™^ mouse. We thank L. Cagnacci for technical support and S. Pellegrino-Corby, M. Kurz and F. Michel from the INMED animal and imaging facilities (InMagic). We are grateful to Dr. S. Reichinnek for valuable scientific input, to Dr. S. Sarno for her help on statistical analysis. A table including all statistical tests performed is included in the supplementary material.

## Supplementary Information

### Recapitulative Table

#### Summary of electrophysiological and morphometric properties of fate-mapped CA1PNs

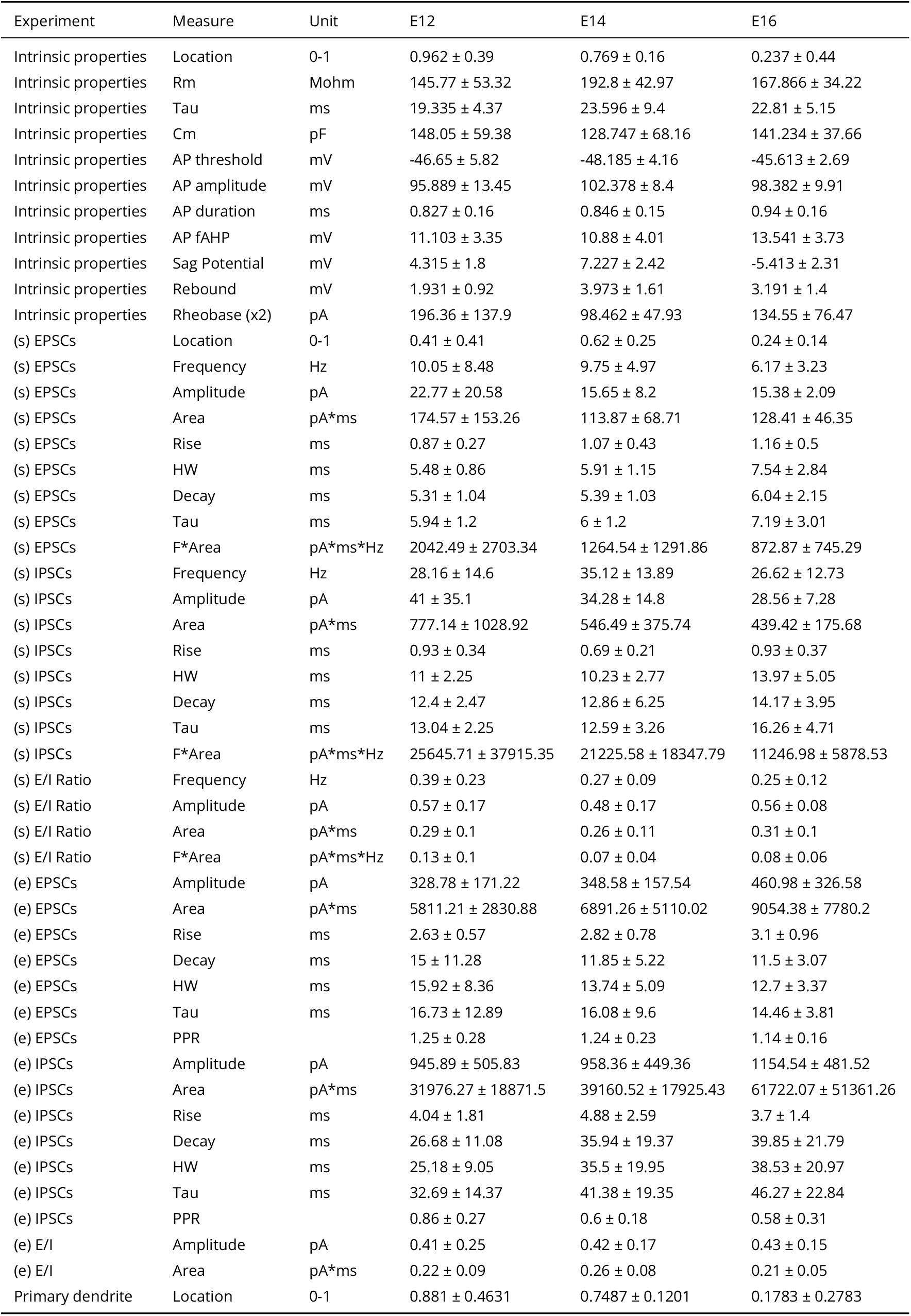
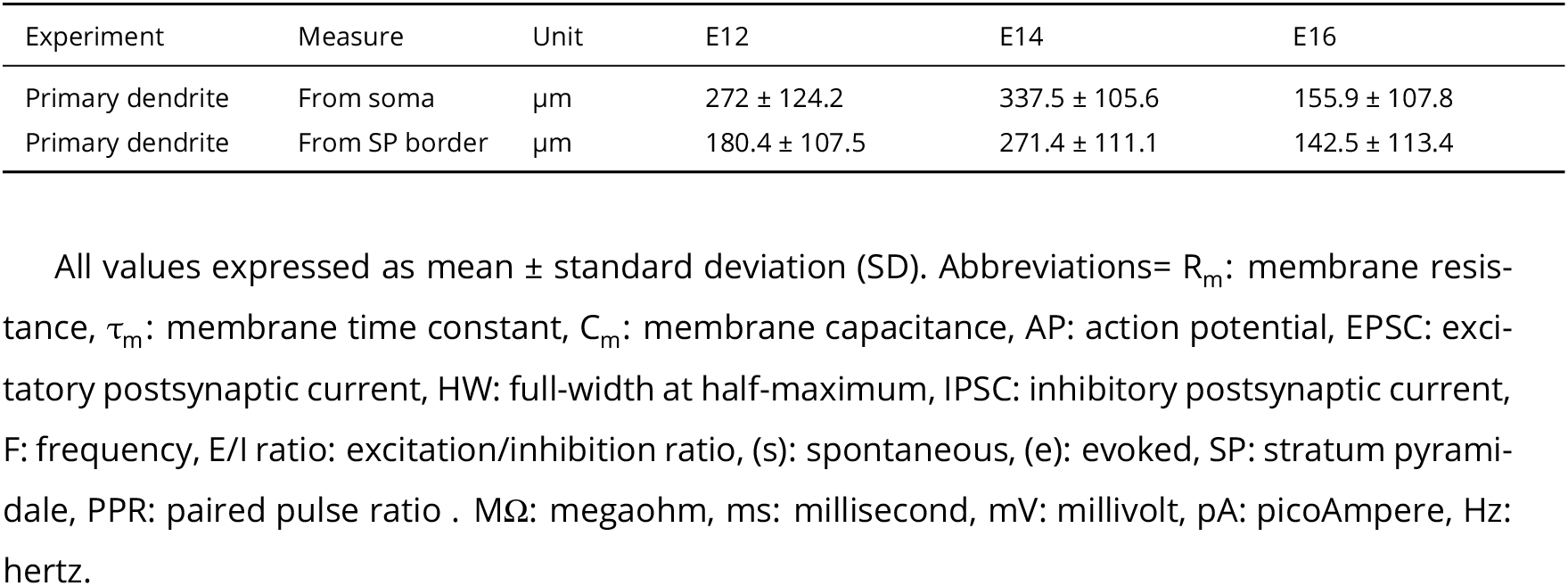

**Summary of electrophysiological and morphometric properties of fate-mapped CA1PNs.**
| Experiment | Measure | Unit | E12 | E14 | E16 |
| --- | --- | --- | --- | --- | --- |
| Intrinsic properties | Location | 0-1 | 0.962 ± 0.39 | 0.769 ± 0.16 | 0.237 ± 0.44 |
| Intrinsic properties | Rm | Mohm | 145.77 ± 53.32 | 192.8 ± 42.97 | 167.866 ± 34.22 |
| Intrinsic properties | Tau | ms | 19.335 ± 4.37 | 23.596 ± 9.4 | 22.81 ± 5.15 |
| Intrinsic properties | Cm | pF | 148.05 ± 59.38 | 128.747 ± 68.16 | 141.234 ± 37.66 |
| Intrinsic properties | AP threshold | mV | -46.65 ± 5.82 | -48.185 ± 4.16 | -45.613 ± 2.69 |
| Intrinsic properties | AP amplitude | mV | 95.889 ± 13.45 | 102.378 ± 8.4 | 98.382 ± 9.91 |
| Intrinsic properties | AP duration | ms | 0.827 ± 0.16 | 0.846 ± 0.15 | 0.94 ± 0.16 |
| Intrinsic properties | AP fAHP | mV | 11.103 ± 3.35 | 10.88 ± 4.01 | 13.541 ± 3.73 |
| Intrinsic properties | Sag Potential | mV | 4.315 ± 1.8 | 7.227 ± 2.42 | -5.413 ± 2.31 |
| Intrinsic properties | Rebound | mV | 1.931 ± 0.92 | 3.973 ± 1.61 | 3.191 ± 1.4 |
| Intrinsic properties | Rheobase (x2) | pA | 196.36 ± 137.9 | 98.462 ± 47.93 | 134.55 ± 76.47 |
| (s) EPSCs | Location | 0-1 | 0.41 ± 0.41 | 0.62 ± 0.25 | 0.24 ± 0.14 |
| (s) EPSCs | Frequency | Hz | 10.05 ± 8.48 | 9.75 ± 4.97 | 6.17 ± 3.23 |
| (s) EPSCs | Amplitude | pA | 22.77 ± 20.58 | 15.65 ± 8.2 | 15.38 ± 2.09 |
| (s) EPSCs | Area | pA*ms | 174.57 ± 153.26 | 113.87 ± 68.71 | 128.41 ± 46.35 |
| (s) EPSCs | Rise | ms | 0.87 ± 0.27 | 1.07 ± 0.43 | 1.16 ± 0.5 |
| (s) EPSCs | HW | ms | 5.48 ± 0.86 | 5.91 ± 1.15 | 7.54 ± 2.84 |
| (s) EPSCs | Decay | ms | 5.31 ± 1.04 | 5.39 ± 1.03 | 6.04 ± 2.15 |
| (s) EPSCs | Tau | ms | 5.94 ± 1.2 | 6 ± 1.2 | 7.19 ± 3.01 |
| (s) EPSCs | F*Area | pA*ms*Hz | 2042.49 ± 2703.34 | 1264.54 ± 1291.86 | 872.87 ± 745.29 |
| (s) IPSCs | Frequency | Hz | 28.16 ± 14.6 | 35.12 ± 13.89 | 26.62 ± 12.73 |
| (s) IPSCs | Amplitude | pA | 41 ± 35.1 | 34.28 ± 14.8 | 28.56 ± 7.28 |
| (s) IPSCs | Area | pA*ms | 777.14 ± 1028.92 | 546.49 ± 375.74 | 439.42 ± 175.68 |
| (s) IPSCs | Rise | ms | 0.93 ± 0.34 | 0.69 ± 0.21 | 0.93 ± 0.37 |
| (s) IPSCs | HW | ms | 11 ± 2.25 | 10.23 ± 2.77 | 13.97 ± 5.05 |
| (s) IPSCs | Decay | ms | 12.4 ± 2.47 | 12.86 ± 6.25 | 14.17 ± 3.95 |
| (s) IPSCs | Tau | ms | 13.04 ± 2.25 | 12.59 ± 3.26 | 16.26 ± 4.71 |
| (s) IPSCs | F*Area | pA*ms*Hz | 25645.71 ± 37915.35 | 21225.58 ± 18347.79 | 11246.98 ± 5878.53 |
| (s) E/I Ratio | Frequency | Hz | 0.39 ± 0.23 | 0.27 ± 0.09 | 0.25 ± 0.12 |
| (s) E/I Ratio | Amplitude | pA | 0.57 ± 0.17 | 0.48 ± 0.17 | 0.56 ± 0.08 |
| (s) E/I Ratio | Area | pA*ms | 0.29 ± 0.1 | 0.26 ± 0.11 | 0.31 ± 0.1 |
| (s) E/I Ratio | F*Area | pA*ms*Hz | 0.13 ± 0.1 | 0.07 ± 0.04 | 0.08 ± 0.06 |
| (e) EPSCs | Amplitude | pA | 328.78 ± 171.22 | 348.58 ± 157.54 | 460.98 ± 326.58 |
| (e) EPSCs | Area | pA*ms | 5811.21 ± 2830.88 | 6891.26 ± 5110.02 | 9054.38 ± 7780.2 |
| (e) EPSCs | Rise | ms | 2.63 ± 0.57 | 2.82 ± 0.78 | 3.1 ± 0.96 |
| (e) EPSCs | Decay | ms | 15 ± 11.28 | 11.85 ± 5.22 | 11.5 ± 3.07 |
| (e) EPSCs | HW | ms | 15.92 ± 8.36 | 13.74 ± 5.09 | 12.7 ± 3.37 |
| (e) EPSCs | Tau | ms | 16.73 ± 12.89 | 16.08 ± 9.6 | 14.46 ± 3.81 |
| (e) EPSCs | PPR |  | 1.25 ± 0.28 | 1.24 ± 0.23 | 1.14 ± 0.16 |
| (e) IPSCs | Amplitude | pA | 945.89 ± 505.83 | 958.36 ± 449.36 | 1154.54 ± 481.52 |
| (e) IPSCs | Area | pA*ms | 31976.27 ± 18871.5 | 39160.52 ± 17925.43 | 61722.07 ± 51361.26 |
| (e) IPSCs | Rise | ms | 4.04 ± 1.81 | 4.88 ± 2.59 | 3.7 ± 1.4 |
| (e) IPSCs | Decay | ms | 26.68 ± 11.08 | 35.94 ± 19.37 | 39.85 ± 21.79 |
| (e) IPSCs | HW | ms | 25.18 ± 9.05 | 35.5 ± 19.95 | 38.53 ± 20.97 |
| (e) IPSCs | Tau | ms | 32.69 ± 14.37 | 41.38 ± 19.35 | 46.27 ± 22.84 |
| (e) IPSCs | PPR |  | 0.86 ± 0.27 | 0.6 ± 0.18 | 0.58 ± 0.31 |
| (e) E/I | Amplitude | pA | 0.41 ± 0.25 | 0.42 ± 0.17 | 0.43 ± 0.15 |
| (e) E/I | Area | pA*ms | 0.22 ± 0.09 | 0.26 ± 0.08 | 0.21 ± 0.05 |
| Primary dendrite | Location | 0-1 | 0.881 ± 0.4631 | 0.7487 ± 0.1201 | 0.1783 ± 0.2783 |
| Primary dendrite | From soma | $\mu\text{m}$ | $272 \pm 124.2$ | $337.5 \pm 105.6$ | $155.9 \pm 107.8$ |
| Primary dendrite | From SP border | $\mu\text{m}$ | $180.4 \pm 107.5$ | $271.4 \pm 111.1$ | $142.5 \pm 113.4$ |

All values expressed as mean ± standard deviation (SD). Abbreviations= $R_m$ : membrane resis- tance, $\tau_m$ : membrane time constant, $C_m$ : membrane capacitance, AP: action potential, EPSC: exci- tatory postsynaptic current, HW: full-width at half-maximum, IPSC: inhibitory postsynaptic current, F: frequency, E/I ratio: excitation/inhibition ratio, (s): spontaneous, (e): evoked, SP: stratum pyrami- dale, PPR: paired pulse ratio . $\text{M}\Omega$ : megaohm, ms: millisecond, mV: millivolt, pA: picoAmpere, Hz: hertz.

**Figure 4–Figure supplement 1.**
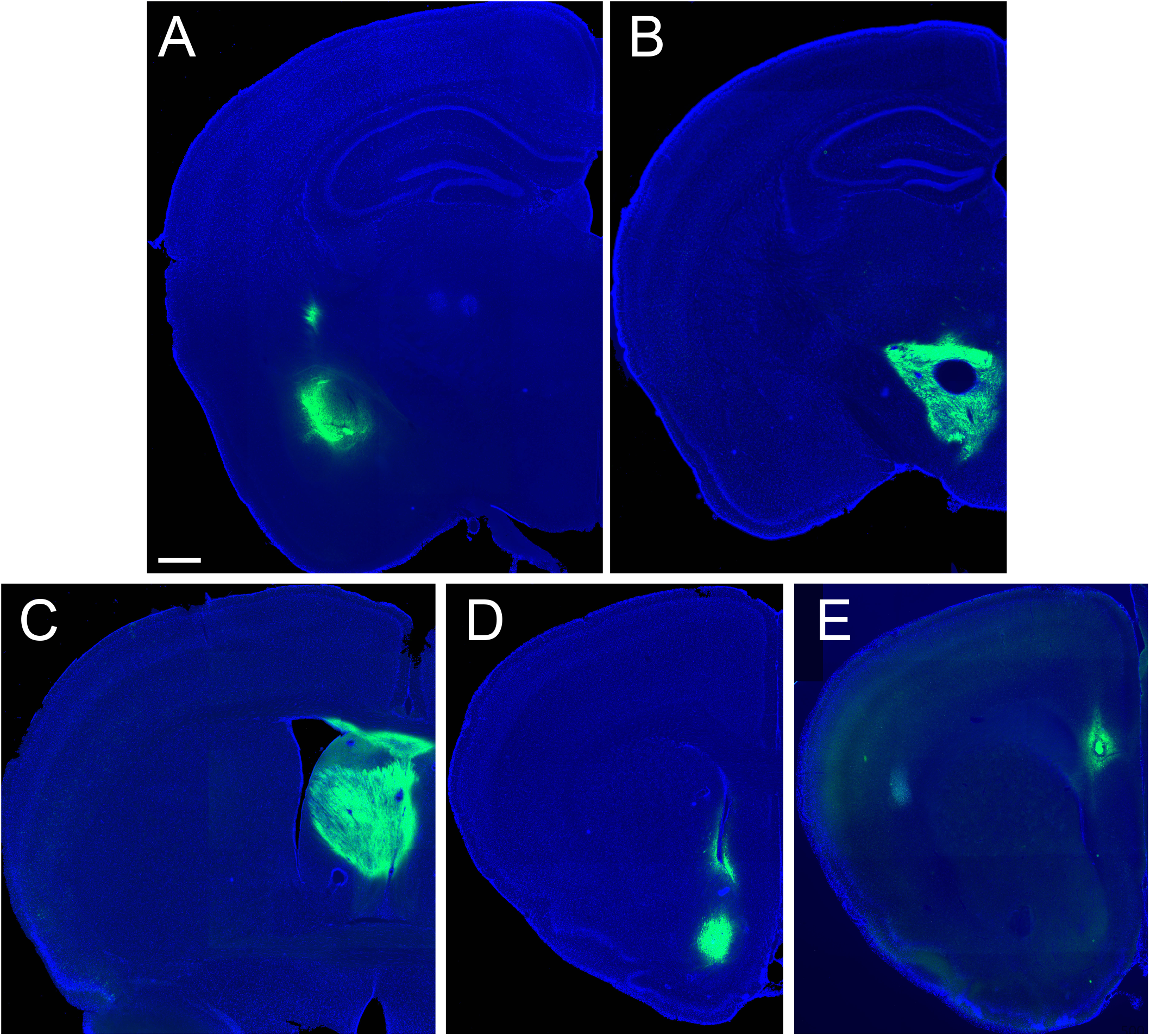
Representative sections including the injection site of Cholera toxin subunit B retrograde tracer, coupled to Alexa647 (here shown in green, DAPI: blue). A: Amygdala; (B) Lateral Hypothalamic Area (LHA); C: Lateral Septum (LS), D: Nucleus Accumbens shell (NAcc); E: medial prefrontal cortex (mPFC). Scalebar: 500 µm.

## References

Allene C, Picardo MA, Becq H, Miyoshi G, Fishell G, Cossart R. Dynamic changes in interneuron morphophysiological properties mark the maturation of hippocampal network activity. Journal of Neuroscience. 2012; 32(19):6688–6698. doi: 10.1523/JNEUROSCI.0081-12.2012.

Angevine JB. Time of neuron origin in the hippocampal region. Experimental neurology Supplement. 1965 10; 11:1–39. http://www.sciencedirect.com/science/article/pii/0014488665901214, doi 10.1016/0014-4886(65)90121-4.

Arszovszki A, Borhegyi Z, Klausberger T. Three axonal projection routes of individual pyramidal cells in the ventral CA1 hippocampus. Frontiers in neuroanatomy. 2014 1; 8:53. http://journal.frontiersin.org/article/10.3389/fnana.2014.00053/full, doi: 10.3389/fnana.2014.00053.

Bayer SA. Development of the hippocampal region in the rat. I. Neurogenesis examined with 3H-thymidine autoradiography. The Journal of comparative neurology. 1980 3; 190(1):87–114. http://www.ncbi.nlm.nih.gov/pubmed/7381056, doi: 10.1002/cne.901900107.

Bittner KC, Milstein AD, Grienberger C, Romani S, Magee JC. Behavioral time scale synaptic plasticity underlies CA1 place fields. Science. 2017; 357(6355). http://science.sciencemag.org.gate2.inist.fr/content/357/6355/1033.full.

Bocchio M, Gouny C, Angulo-Garcia D, Toulat T, Tressard T, Quiroli E, Baude A, Cossart R. Hippocampal hub neurons maintain distinct connectivity throughout their lifetime. Nature Communications. 2020; 11(1). http://dx.doi.org/10.1038/s41467-020-18432-6, doi:10.1038/s41467-020-18432-6.

Carpenter AE, Jones TR, Lamprecht MR, Clarke C, Kang IH, Friman O, Guertin DA, Chang JH, Lindquist RA, Moffat J, Golland P, Sabatini DM. CellProfiler: Image analysis software for identifying and quantifying cell phenotypes. Genome Biology. 2006 10; 7(10):R100. http://genomebiology.biomedcentral.com/articles/10.1186/gb-2006-7-10-r100, doi: 10.1186/gb-2006-7-10-r100.

Caviness VS. Time of neuron origin in the hippocampus and dentate gyrus of normal and reeler mutant mice: an autoradiographic analysis. The Journal of comparative neurology. 1973 9; 151(2):113–20. http://www.ncbi.nlm.nih.gov/pubmed/4744470, doi: 10.1002/cne.901510203.

Cembrowski MS, Bachman JL, Wang L, Sugino K, Shields BC, Spruston N. Spatial Gene-Expression Gradients Underlie Prominent Heterogeneity of CA1 Pyramidal Neurons. Neuron. 2016 1; 89(2):351–368. http://www.sciencedirect.com/science/article/pii/S0896627315010867, doi: 10.1016/j.neuron.2015.12.013.

Cenquizca LA, Swanson LW. Spatial organization of direct hippocampal field CA1 axonal projections to the rest of the cerebral cortex. Brain Research Reviews. 2007 11; 56(1):1–26. http://www.sciencedirect.com/science/article/pii/S0165017307000732?via%3Dihub, doi: 10.1016/j.brainresrev.2007.05.002.

Ciocchi S, Passecker J, Malagon-Vina H, Mikus N, Klausberger T. Selective information routing by ventral hippocampal CA1 projection neurons. Science. 2015 4; 348(6234):560–563. http://www.sciencemag.org.gate2.inist.fr/content/348/6234/560.full, doi: 10.1126/science.aaa3245.

Cossart R, Hirsch JC, Cannon RC, Dinoncourt C, Wheal HV, Ben-Ari Y, Esclapez M, Bernard C. Distribution of spontaneous currents along the somato-dendritic axis of rat hippocampal CA1 pyramidal neurons. Neuroscience. 2000; doi: 10.1016/S0306-4522(00)00231-1.

D’Amour JA, Ekins TG, Ghanatra S, Yuan X, McBain CJ, Ganatra S, Yuan X, McBain CJ. Aberrant sorting of hippocampal complex pyramidal cells in Type I Lissencephaly alters topological innervation. eLife. 2020 6; 9:1–26. doi: 10.1101/2020.02.05.935775.

Danielson N, Zaremba J, Kaifosh P, Bowler J, Ladow M, Losonczy A. Sublayer-Specific Coding Dynamics during Spatial Navigation and Learning in Hippocampal Area CA1. Neuron. 2016; 91(3):652–665. doi: 10.1016/j.neuron.2016.06.020.

Deguchi Y, Donato F, Galimberti I, Cabuy E, Caroni P. Temporally matched subpopulations of selectively interconnected principal neurons in the hippocampus. Nature neuroscience. 2011 4; 14(4):495–504. http://www.nature.com.gate2.inist.fr/neuro/journal/v14/n4/full/nn.2768.html, doi: 10.1038/nn.2768.

Donato F, Jacobsen RI, Moser MB, Moser EI. Stellate cells drive maturation of the entorhinal-hippocampal circuit. Science. 2017 2; 355(6330):eaai8178. http://www.ncbi.nlm.nih.gov/pubmed/28154241 http://www.sciencemag.org/lookup/doi/10.1126/science.aai8178, doi: 10.1126/science.aai8178.

Dougherty KA. Differential developmental refinement of the intrinsic electrophysiological properties of CA1 pyramidal neurons from the rat dorsal and ventral hippocampus. Hippocampus. 2019 9; p. hipo.23152. https://onlinelibrary.wiley.com/doi/abs/10.1002/hipo.23152, doi: 10.1002/hipo.23152.

Dougherty KA, Islam T, Johnston D. Intrinsic excitability of CA1 pyramidal neurones from the rat dorsal and ventral hippocampus. The Journal of Physiology. 2012 11; 590(22):5707–5722. http://doi.wiley.com/10.1113/jphysiol.2012.242693, doi: 10.1113/jphysiol.2012.242693.

Epsztein J, Brecht M, Lee A. Intracellular Determinants of Hippocampal CA1 Place and Silent Cell Activity in a Novel Environment. Neuron. 2011; 70(1):109–120. doi: 10.1016/j.neuron.2011.03.006.

Fattahi M, Sharif F, Geiller T, Royer S. Differential representation of landmark and self-motion information along the CA1 radial axis: Self-motion generated place fields shift toward landmarks during septal inactivation. Journal of Neuroscience. 2018 7; 38(30):6766–6778. https://www.jneurosci.org/content/38/30/6766 https://www.jneurosci.org/content/38/30/6766.abstract, doi: 10.1523/JNEUROSCI.3211-17.2018.

Franco SJ, Gil-Sanz C, Martinez-Garay I, Espinosa A, Harkins-Perry SR, Ramos C, Müller U. Fate-restricted neural progenitors in the mammalian cerebral cortex. Science. 2012; doi: 10.1126/science.1223616.

Geiller T, Fattahi M, Choi JS, Royer S. Place cells are more strongly tied to landmarks in deep than in superficial CA1. Nature Communications. 2017 2; 8:14531. http://www.nature.com/doifinder/10.1038/ncomms14531, doi: 10.1038/ncomms14531.

Gergues MM, Han KJ, Choi HS, Brown B, Clausing KJ, Turner VS, Vainchtein ID, Molofsky AV, Kheirbek MA. Circuit and molecular architecture of a ventral hippocampal network. Nature Neuroscience. 2020; http://dx.doi.org/10.1038/s41593-020-0705-8, doi: 10.1038/s41593-020-0705-8.

Graves AR, Moore SJ, Bloss EB, Mensh BD, Kath WL, Spruston N. Hippocampal pyramidal neurons comprise two distinct cell types that are countermodulated by metabotropic receptors. Neuron. 2012 11; 76(4):776–89. http://www.cell.com/article/S0896627312008902/fulltext, doi: 10.1016/j.neuron.2012.09.036.

van Groen T, Wyss JM. Extrinsic projections from area CA1 of the rat hippocampus: olfactory, cortical, subcortical, and bilateral hippocampal formation projections. The Journal of comparative neurology. 1990 12; 302(3):515–28. http://www.ncbi.nlm.nih.gov/pubmed/1702115, doi: 10.1002/cne.903020308.

Gulyás AI, Tóth K, McBain CJ, Freund TF. Stratum radiatum giant cells: A type of principal cell in the rat hippocampus. European Journal of Neuroscience. 1998; 10(12):3813–3822. doi: 10.1046/j.1460-9568.1998.00402.x.

Gupta A, Elgammal FS, Proddutur A, Shah S, Santhakumar V. Decrease in tonic inhibition contributes to increase in dentate semilunar granule cell excitability after brain injury. Journal of Neuroscience. 2012; doi: 10.1523/JNEUROSCI.4141-11.2012.

Hartzell AL, Burke SN, Hoang LT, Lister JP, Rodriguez CN, Barnes CA. Transcription of the immediate-early gene Arc in CA1 of the hippocampus Reveals activity differences along the proximodistal axis that are Attenuated by advanced age. Journal of Neuroscience. 2013 2; 33(8):3424–3433. https://www.jneurosci.org/content/33/8/3424https://www.jneurosci.org/content/33/8/3424.abstract, doi: 10.1523/JNEUROSCI.4727-12.2013.

Henriksen EJ, Colgin LL, Barnes CA, Witter MP, Moser MB, Moser EI. Spatial Representation along the Proximodistal Axis of CA1. Neuron. 2010 10; 68(1):127–137. https://www.sciencedirect.com/science/article/pii/S0896627310006859, doi: 10.1016/J.NEURON.2010.08.042.

Jabaudon D, Fate and freedom in developing neocortical circuits; 2017. doi: 10.1038/ncomms16042.

Jarsky T, Mady R, Kennedy B, Spruston N. Distribution of bursting neurons in the CA1 region and the subiculum of the rat hippocampus. The Journal of comparative neurology. 2008 2; 506(4):535–47. http://www.ncbi.nlm.nih.gov/pubmed/18067146, doi: 10.1002/cne.21564.

Jimenez JC, Su K, Goldberg AR, Luna VM, Biane JS, Ordek G, Zhou P, Ong SK, Wright MA, Zweifel L, Paninski L, Hen R, Kheirbek MA. Anxiety Cells in a Hippocampal-Hypothalamic Circuit. Neuron. 2018 2; 97(3):670–683. http://www.ncbi.nlm.nih.gov/pubmed/29397273, doi: 10.1016/j.neuron.2018.01.016.

Jin J, Maren S. Fear renewal preferentially activates ventral hippocampal neurons projecting to both amygdala and prefrontal cortex in rats. Scientific Reports. 2015; 5:8388. http://www.pubmedcentral.nih.gov/articlerender.fcgi?artid=4323647&tool=pmcentrez&rendertype=abstract, doi: 10.1038/srep08388.

Kaczorowski CC, Disterhoft J, Spruston N. Stability and plasticity of intrinsic membrane properties in hippocampal CA1 pyramidal neurons: Effects of internal anions. The Journal of Physiology. 2007 2; 578(3):799–818. http://www.ncbi.nlm.nih.gov/pubmed/17138601http://www.pubmedcentral.nih.gov/articlerender.fcgi?artid=PMC2151348http://doi.wiley.com/10.1113/jphysiol.2006.124586, doi: 10.1113/jphysiol.2006.124586.

Kim WB, Cho JH. Synaptic Targeting of Double-Projecting Ventral CA1 Hippocampal Neurons to the Medial Prefrontal Cortex and Basal Amygdala. The Journal of Neuroscience. 2017; 37(19):4868–4882. http://www.jneurosci.org/lookup/doi/10.1523/JNEUROSCI.3579-16.2017, doi: 10.1523/JNEUROSCI.3579-16.2017.

Kitanishi T, Ikegaya Y, Matsuki N, Yamada MK. Experience-dependent, rapid structural changes in hippocampal pyramidal cell spines. Cerebral Cortex. 2009; doi: 10.1093/cercor/bhp012.

Kohara K, Pignatelli M, Rivest AJ, Jung HY, Kitamura T, Suh J, Frank D, Kajikawa K, Mise N, Obata Y, Wickersham IR, Tonegawa S. Cell type-specific genetic and optogenetic tools reveal hippocampal CA2 circuits. Nature neuroscience. 2014 2; 17(2):269–79. http://www.nature.com.gate2.inist.fr/neuro/journal/v17/n2/full/nn.3614.html, doi: 10.1038/nn.3614.

Lee SH, Marchionni I, Bezaire M, Varga C, Danielson N, Lovett-Barron M, Losonczy A, Soltesz I. Parvalbumin-positive basket cells differentiate among hippocampal pyramidal cells. Neuron. 2014 6; 82(5):1129–44. http://www.sciencedirect.com/science/article/pii/S0896627314003365, doi: 10.1016/j.neuron.2014.03.034.

Li Y, Xu J, Liu Y, Zhu J, Liu N, Zeng W, Huang N, Rasch MJ, Jiang H, Gu X, Li X, Luo M, Li C, Teng J, Chen J, Zeng S, Lin L, Zhang X. A distinct entorhinal cortex to hippocampal CA1 direct circuit for olfactory associative learning. Nature Neuroscience. 2017 4; 20(4):559–570. http://www.nature.com/articles/nn.4517, doi: 10.1038/nn.4517.

Lodato S, Shetty AS, Arlotta P, Cerebral cortex assembly: Generating and reprogramming projection neuron diversity; 2015. doi: 10.1016/j.tins.2014.11.003.

Madisen L, Zwingman TA, Sunkin SM, Oh SW, Zariwala HA, Gu H, Ng LL, Palmiter RD, Hawrylycz MJ, Jones AR, Lein ES, Zeng H. A robust and high-throughput Cre reporting and characterization system for the whole mouse brain. Nature Neuroscience. 2010; doi: 10.1038/nn.2467.

Marissal T, Bonifazi P, Picardo MA, Nardou R, Petit LF, Baude A, Fishell GJ, Ben-Ari Y, Cossart R. Pioneer gluta-matergic cells develop into a morpho-functionally distinct population in the juvenile CA3 hippocampus. Nature communications. 2012 1; 3:1316. http://www.pubmedcentral.nih.gov/articlerender.fcgi?artid=3535425&tool=pmcentrez&rendertype=abstract, doi: 10.1038/ncomms2318.

Maroso M, Szabo GG, Kim HK, Alexander A, Bui AD, Lee SH, Lutz B, Soltesz I. Cannabinoid Control of Learning and Memory through HCN Channels. Neuron. 2016; 89(5):1059–1073. http://www.sciencedirect.com/science/article/pii/S0896627316000489, doi: 10.1016/j.neuron.2016.01.023.

Masurkar AV, Srinivas KV, Brann DH, Warren R, Lowes DC, Siegelbaum SA. Medial and Lateral Entorhinal Cortex Differentially Excite Deep versus Superficial CA1 Pyramidal Neurons. CellReports. 2017; 18:148–160. http://dx.doi.org/10.1016/j.celrep.2016.12.012, doi: 10.1016/j.celrep.2016.12.012.

Masurkar AV, Tian C, Warren R, Reyes I, Lowes DC, Brann DH, Siegelbaum SA. Postsynaptic integrative properties of dorsal CA1 pyramidal neuron subpopulations. Journal of Neurophysiology. 2020 1; doi: 10.1152/jn.00397.2019.

Mathews EA, Morgenstern NA, Piatti VC, Zhao C, Jessberger S, Schinder AF, Gage FH. A distinctive layering pattern of mouse dentate granule cells is generated by developmental and adult neurogenesis. Journal of Comparative Neurology. 2010; doi: 10.1002/cne.22489.

Mizuseki K, Diba K, Pastalkova E, Buzsáki G. Hippocampal CA1 pyramidal cells form functionally distinct sublayers. Nature neuroscience. 2011 9; 14(9):1174–. http://www.nature.com/neuro/journal/v14/n9/abs/nn.2894.html, doi: 10.1038/nn.2894.Hippocampal.

Nasrallah K, Therreau L, Robert V, Huang AJY, McHugh TJ, Piskorowski RA, Chevaleyre V. Routing Hippocampal Information Flow through Parvalbumin Interneuron Plasticity in Area CA2. Cell Reports. 2019; 27(1):86–98. https://doi.org/10.1016/j.celrep.2019.03.014, doi: 10.1016/j.celrep.2019.03.014.

Okuyama T, Kitamura T, Roy DS, Itohara S, Tonegawa S. Ventral CA1 neurons store social memory. Science (New York, NY). 2016 9; 353(6307):1536–1541. http://www.ncbi.nlm.nih.gov/pubmed/27708103http://www.pubmedcentral.nih.gov/articlerender.fcgi?artid=PMC5493325, doi: 10.1126/science.aaf7003.

Oliva A, Fernández-Ruiz A, Buzsáki G, Berényi A. Spatial coding and physiological properties of hip-pocampal neurons in the Cornu Ammonis subregions. Hippocampus. 2016; 26(12):1593–1607. doi: 10.1002/hipo.22659.

Parfitt GM, Nguyen R, Bang JY, Aqrabawi AJ, Tran MM, Seo DK, Richards BA, Kim JC. Bidirectional Control of Anxiety-Related Behaviors in Mice: Role of Inputs Arising from the Ventral Hippocampus to the Lateral Septum and Medial Prefrontal Cortex. Neuropsychopharmacology. 2017; doi: 10.1038/npp.2017.56.

Pouille F, Scanziani M. Routing of spike series by dynamic circuits in the hippocampus. Nature. 2004; doi: 10.1038/nature02615.

Save L, Baude A, Cossart R. Temporal Embryonic Origin Critically Determines Cellular Physiology in the Dentate Gyrus. Cerebral Cortex. 2019; 29(6):2639–2652. doi: 10.1093/cercor/bhy132.

Sharif F, Tayebi B, Buzsáki G, Royer S, Fernandez-Ruiz A. Subcircuits of Deep and Superficial CA1 Place Cells Support Effcient Spatial Coding across Heterogeneous Environments. Neuron. 2020; p. 1–14. doi: 10.1016/j.neuron.2020.10.034.

Soltesz I, Losonczy A. CA1 pyramidal cell diversity enabling parallel information processing in the hippocampus. Nature Neuroscience. 2018 3; 21(4):484–493. https://www-nature-com.gate2.inist.fr/articles/s41593-018-0118-0?WT.ec_id=NEURO-201804&spMailingID=56319302&spUserID=MjEyMDk1MTIwNjM1S0&spJobID=1380254163&spReportId=MTM4MDI1NDE2MwS2, doi: 10.1038/s41593-018-0118-0.

Thomson AM, Bannister AP, Hughes DI, Pawelzik H. Differential sensitivity to Zolpidem of IPSPs activated by morphologically identified CA1 interneurons in slices of rat hippocampus. European Journal of Neuroscience. 2000; doi: 10.1046/j.1460-9568.2000.00915.x.

Valero M, Cid E, Averkin RG, Aguilar J, Sanchez-Aguilera A, Viney TJ, Gomez-Dominguez D, Bellistri E, de la Prida LM. Determinants of different deep and superficial CA1 pyramidal cell dynamics during sharp-wave ripples. Nature neuroscience. 2015 9; 18(9):1281–90. http://dx.doi.org/10.1038/nn.4074, doi: 10.1038/nn.4074.

Valero M, de la Prida LM. The hippocampus in depth: a sublayer-specific perspective of entorhinal–hippocampal function. Current Opinion in Neurobiology. 2018 10; 52:107–114. https://www-sciencedirect-com.gate2.inist.fr/science/article/pii/S0959438817302994?via%3Dihub, doi: 10.1016/j.conb.2018.04.013.

West AE, Griffth EC, Greenberg ME. Regulation of transcription factors by neuronal activity. Nature Reviews Neuroscience. 2002; doi: 10.1038/nrn987.

Xu C, Krabbe S, Gründemann J, Botta P, Fadok JP, Osakada F, Saur D, Grewe BF, Schnitzer MJ, Callaway EM, Lüthi A. Distinct Hippocampal Pathways Mediate Dissociable Roles of Context in Memory Retrieval. Cell. 2016; 167(4):961–972. doi: 10.1016/j.cell.2016.09.051.

Xu HT, Han Z, Gao P, He S, Li Z, Shi W, Kodish O, Shao W, Brown KN, Huang K, Shi SH. Distinct lineage-dependent structural and functional organization of the hippocampus. Cell. 2014 6; 157(7):1552–64. http://www.sciencedirect.com/science/article/pii/S0092867414006102, doi: 10.1016/j.cell.2014.03.067.

